# Correlative mapping links extracellular matrix architecture to local mechanics in cardiac aging

**DOI:** 10.64898/2026.11.21.768346

**Authors:** Md. Faris H. Ramli, Alokendra Ghosh, Ranmadusha Hengst, Avery Rui Sun, Huan Ting Ong, Yuting Lou, Hui Ting Ong, Murat Shagirov, Xingyu Shen, Kevin Nicholas, Gianluca Grenci, Tetsuya Hiraiwa, Jennifer L. Young

## Abstract

Cardiac aging is accompanied by extracellular matrix (ECM) remodeling, but how changes in matrix architecture relate to local tissue mechanics remains poorly understood. Here, we quantified six image-derived architectural descriptors across selected ECM components in young and aged murine myocardium and integrated them into an architectural index that we term ‘ArchI’. Age-associated architectural changes differed among ECM components, with higher ArchI values observed for fibrillar collagen, fibronectin, and laminin-associated networks. To examine how defined structural changes affect the component descriptors, we developed controlled synthetic geometries and a stochastic network model. These analyses revealed index sensitivity to matrix occupancy, fiber width, and network continuity. We then combined nanoindentation of fresh cardiac tissue slices with subsequent immunofluorescence staining and imaging of collagen to correlate local tissue stiffness with ECM architecture in corresponding regions. Aged myocardium exhibited higher ArchI and Young’s modulus, with local heterogeneity in their spatial correspondence. Age-adjusted regression revealed a positive association between ArchI and Young’s modulus, with a steeper association in aged myocardium. Together, these findings reveal component-specific ECM remodeling during aging and provide a means to examine spatial structure-mechanics relationships in the aging heart.

## Introduction

Age-associated structural and mechanical remodeling can impair cardiac function before overt cardiovascular disease. Thus, defining matrix changes that accompany this remodeling may help explain how tissue homeostasis is altered with age [1–5]. Unlike pathological remodeling, such as replacement fibrosis following myocardial infarction, cardiac aging can involve matrix accumulation and modification [6,7]. The myocardium maintains an ordered, anisotropic architecture that supports coordinated contraction and directional force transmission. This complex microenvironment is sustained by the ECM, a highly organized structural network that preserves cardiomyocyte alignment, maintains tissue integrity, and regulates local mechanical signaling [8,9]. Although ECM remodeling is essential for normal tissue homeostasis, aging is associated with changes in matrix composition and architecture [7,10–12]. Such changes may increase tissue stiffness, disrupt local force transmission, and impair the mechanical coordination required for optimal cardiac function. Defining how age-related changes in ECM architecture relate to local tissue stiffness is therefore important for understanding how matrix remodeling contributes to cardiac dysfunction.

The myocardial ECM consists of structurally distinct matrix networks, including interstitial fibrillar collagens and fibronectin, basement membranes containing laminins and collagen IV, and linkers of interstitial and pericellular matrix compartments including collagen VI microfibrils [13,14]. During aging, changes in ECM abundance, crosslinking, and organization may contribute to myocardial stiffening and altered tissue mechanics [6,10]. Multiscale analyses of failing hearts have linked ECM remodeling to changes in matrix organization and mechanics, while studies on engineered myocardial microtissues show that matrix architecture and mechanics influence myofibril organization and contractility [15,16]. These observations highlight the importance of examining age-associated remodeling across individual ECM components.

Conventional assessments of cardiac fibrosis and myocardial stiffening often quantify collagen accumulation or bulk tissue mechanical properties, which do not resolve how local ECM architecture relates to stiffness within individual regions of the heart [15,17,18]. Image analysis tools such as TWOMBLI quantify multiple features of ECM organization [19], and correlative methods such as STIFMap use collagen staining and local mechanical measurements to predict stromal stiffness in breast tumors [20]. However, how architectural remodeling differs among ECM components with age, as well as how these architectures relate directly to local tissue stiffness remains limited.

Here, we investigated age-associated differences in localized ECM architecture and tissue mechanics within the myocardium of young (∼1 month) and aged (18-24 months) mice. We asked whether architectural changes differed among ECM components and whether local fibrillar collagen architecture was correlated with tissue stiffness as a function of age. To address these questions, we first quantified age-associated architectural remodeling across distinct ECM components using multidimensional geometric descriptors derived from ECM imaging. These structural features were subsequently integrated into a composite Architecture Index (ArchI) to identify differences in matrix organization. We then spatially correlated microscale stiffness maps from nanoindentation on fresh cardiac tissue with subsequent ECM imaging using a customized grid-based workflow, which allowed for the relocation of indented tissue regions to examine structure-mechanics relationships. Defining how localized ECM architecture associates with tissue stiffness may help identify structural features to incorporate into biomaterial platforms and experimental models that better recapitulate the aged cardiac microenvironment.

## Results

### Cardiac ECM components undergo distinct age-associated remodeling

Our previous work identified age-associated differences in cardiac ECM composition and collagen architecture [7]. Here, we examined how architectural remodeling differs across ECM components of interstitial, pericellular, and basement membrane networks. To do this, we immunostained fibrillar collagen, fibronectin, collagen VI, laminin, and collagen IV in young and aged myocardial cryosections (**Fig. 1a**). Differences in tissue and ECM network organization were observed between young and aged tissues. Fluorescence intensity analysis revealed age-associated increases in collagen IV and a decrease in laminin, while fibrillar collagen showed a nonsignificant trend toward increased intensity after FDR correction. (**Fig. 1b; Supp. Table S1**). Although fluorescence intensity provides a measure of relative ECM abundance, it does not describe how the matrix is spatially organized. We therefore characterized ECM architecture using six geometric descriptors derived from image masks (**Fig. S1a,b; Supp. Table S6**).

**Figure 1.**
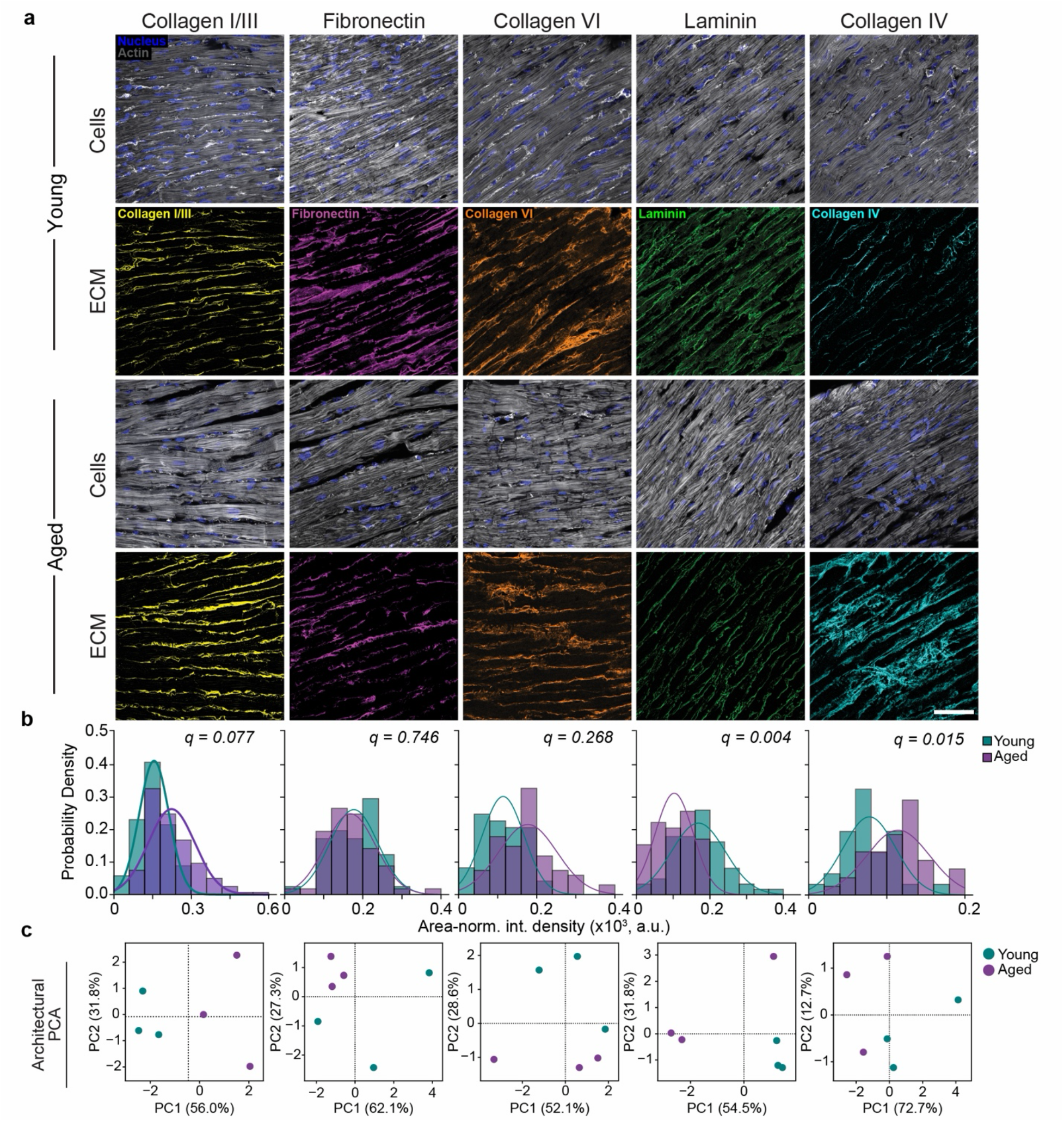
Quantitative characterization of cardiac extracellular matrix architecture identifies age-specific features. (**a**) Immunofluorescence images of cryosectioned young and aged murine cardiac tissue showing the major structural extracellular matrix (ECM) compartments. The interstitial matrix is visualized by fibrillar collagen (yellow) and fibronectin (magenta), the pericellular matrix by collagen VI (orange), and the basement membrane by laminin (green) and collagen IV (cyan). Nuclei are stained with Hoechst (blue), and F-actin is visualized with phalloidin (gray). Scale bar: 50 µm. (**b**) Probability distributions of area-normalized integrated density in young (green) and aged (purple) myocardium. For visualization, the minimum observed intensity for each ECM component was subtracted from both age groups. (**c**) Principal component analysis (PCA) of ECM architectural features from individual biological replicates. Age-group comparisons in (b) were performed using ordinary least squares (OLS) regression with CR1 cluster-robust standard errors clustered by biological replicate (heart). P values were adjusted across ECM components using the Benjamini–Hochberg false discovery rate (FDR) procedure.

These six descriptors represent matrix occupancy (high-density matrix, HDM), space-filling (fractal dimension, FD), variation in matrix-free gaps (lacunarity), apparent fiber thickness (fiber width), fiber orientation dispersion (fiber deviation), and network continuity (hyphal growth unit, HGU). Analysis of the individual descriptors identified component-specific architectural changes with age (**Supp. Table S7**). In aged tissue, fibrillar collagen showed higher HDM, greater apparent fiber width, and lower FD. Fibronectin showed a trend toward greater fiber deviation as opposed to collagen VI. Laminin networks exhibited significantly lower HGU, while collagen IV showed increased apparent fiber width.

Because these changes were distributed across different geometric features, we next used principal component analysis (PCA) to examine whether broader architectural patterns emerged when all six descriptors were considered collectively. PCA performed separately for each ECM component showed varying degrees of separation between young and aged tissues in the first two principal components (**Fig. 1c**). Pairwise correlations also showed that the strength and direction of relationships among descriptors varied across ECM components (**Fig. S1c**). Overall, these analyses identified distinct patterns of age-associated architectural remodeling among ECM components. Because the component-specific analyses represented ECM architecture on separate multivariate axes, we next integrated the geometric descriptors across ECM components to obtain a shared quantitative representation of architectural variation.

### An integrated Architecture Index (ArchI) captures age-associated differences in cardiac ECM organization

To examine ECM architecture across components within a common multivariate space, we performed an integrated PCA incorporating the six standardized geometric descriptors quantified for each ECM component. Young and aged samples separated primarily along the first principal component (PC1) (**Fig. 2a**), which accounted for 48.6% of the total variance (**Supp. Fig. S2b**). We used the corresponding PC1 loadings to derive the Architecture Index (ArchI) as a weighted linear combination of the standardized geometric descriptors (**Supp. Table S4**), providing a single score that summarizes the dominant architectural variation in the dataset. The score was oriented so that higher values corresponded to the age-associated architectural pattern observed in the reference dataset (see Methods section: Architecture Index (ArchI) derivation).

**Figure 2.**
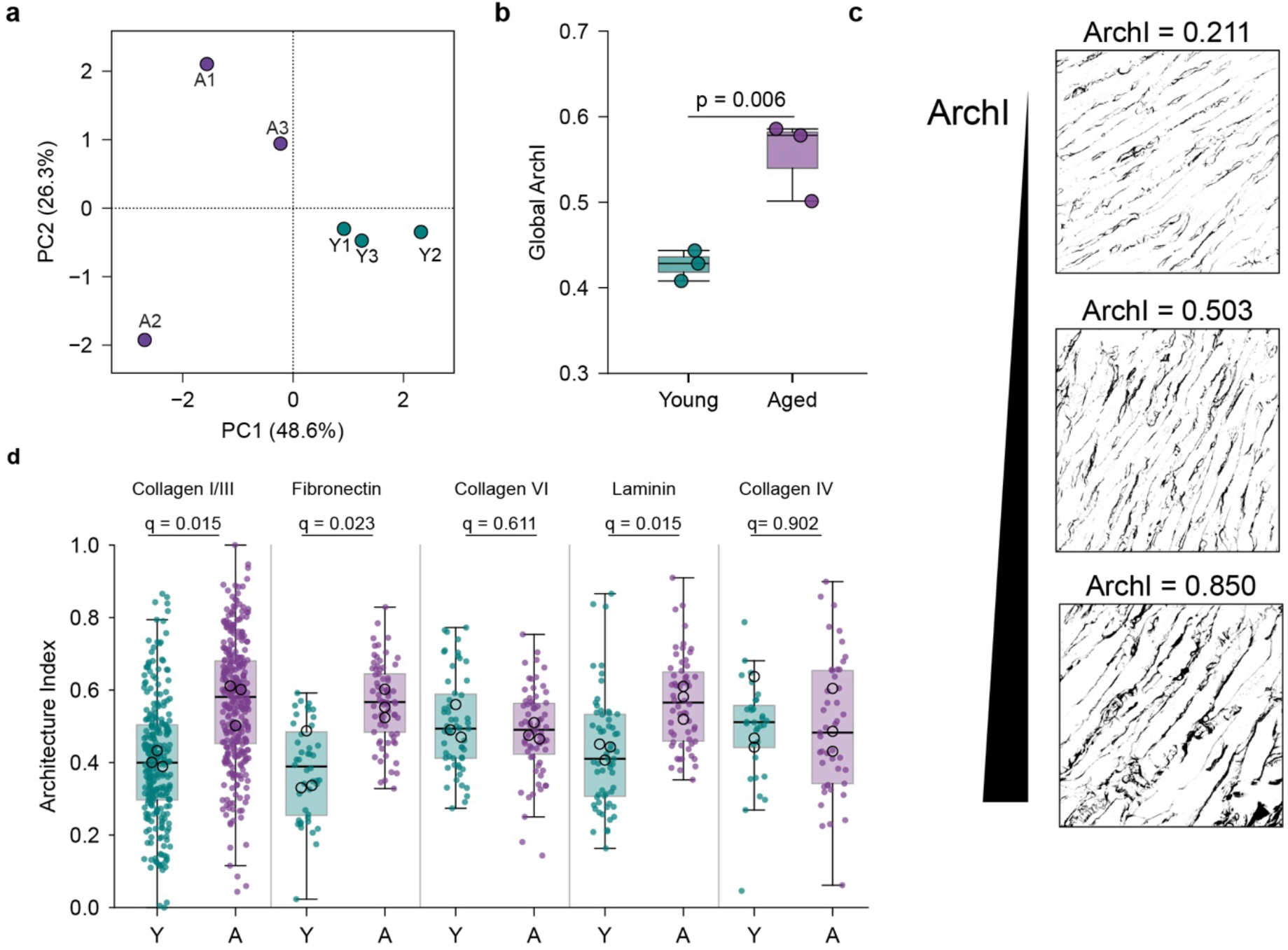
An Architecture Index (ArchI) captures age-associated remodeling across ECM components. **(a)** Principal component analysis (PCA) of geometric descriptors integrated across all ECM components in young (Y1–Y3) and aged (A1–A3) myocardium. **(b)** Architecture Index (ArchI) derived from the integrated PCA, comparing young and aged myocardium. **(c)** Representative ECM architecture images illustrating progressively increasing ArchI values (0.211, 0.503, and 0.850), from lower to higher architectural remodeling. **(d)** Component-specific ArchI distributions for collagen I/III, fibronectin, collagen VI, laminin, and collagen IV in young and aged myocardium. For (b), points represent individual heart-level means. For (d), colored points represent individual regions of interest (ROIs), while open black circles indicate the mean for each biological replicate (heart). In both panels, boxes indicate the interquartile range (IQR), central lines indicate the median, and whiskers extend to the most extreme observations within 1.5 × IQR below the first quartile and above the third quartile. Age-group comparisons were performed using ordinary least squares (OLS) regression with CR1 cluster-robust standard errors clustered by biological replicate (heart). Exact P values are indicated in (b), while P values in (d) were adjusted for multiple comparisons using the Benjamini–Hochberg false discovery rate (FDR) procedure (n = 3 hearts per age group).

Across the analyzed ECM components, aged tissues exhibited significantly higher ArchI values than young tissues (mean values of 0.55 and 0.42, respectively; **Fig. 2b**; *p = 0.0056*; **Supp. Table S2**). Example ECM images with ArchI values spanning ∼0.2-0.9 illustrate the architectural variation summarized by the index (**Fig. 2c**). We next stratified ArchI values by ECM component to examine which matrix networks contributed to this overall difference (**Fig. 2d**). Significant increases were observed for fibrillar collagen, fibronectin, and laminin in aged myocardium (FDR-adjusted *q* = 0.015, 0.023, and 0.015, respectively), whereas no significant differences were detected for collagen VI or collagen IV (**Supp. Table S2**). We next examined how individual structural features contribute to variation in ArchI using experimental descriptor profiles and controlled synthetic geometries.

### Synthetic ECM geometries reveal how structural remodeling alters ArchI

To assess which structural features contribute to variation in ArchI, we examined the geometric descriptor profiles of experimental images spanning the ArchI range (**Fig. 3a; Supp. Fig. S4a**). Within these examples, higher ArchI scores were generally accompanied by lower HGU and greater fiber width. The weighted PC1 contributions illustrate how the six descriptors combine to generate the index (**Fig. 3b**). To evaluate how defined structural changes affect these descriptors, we generated synthetic ECM structures with known geometries and quantified them using a separate implementation of the six descriptor measurements (**Supp. Fig. S3a**). At the heart level, aged tissue was characterized by higher HDM and fiber width and lower FD and HGU (**Fig. 3d**, top row). Controlled perturbation of individual geometric parameters using paired random seeds showed that fiber thickening increased HDM and fiber width, whereas fragmentation reduced HGU. Increasing parallel spacing while widening fibers to maintain constant matrix area increased lacunarity and apparent fiber width (**Fig. 3d; Supp. Fig. S3**). These individual perturbations showed partial directional agreement with the experimental aged-minus-young descriptor pattern, with cosine similarities of 0.47 for thickening, 0.60 for fragmentation, and 0.57 for matched-area spacing (**Supp. Fig. S3**). Here, cosine similarity describes agreement in the direction of the six-descriptor changes, with a value of 1 indicating matching directions. However, none of these perturbations reproduced the complete experimental pattern.

**Figure 3.**
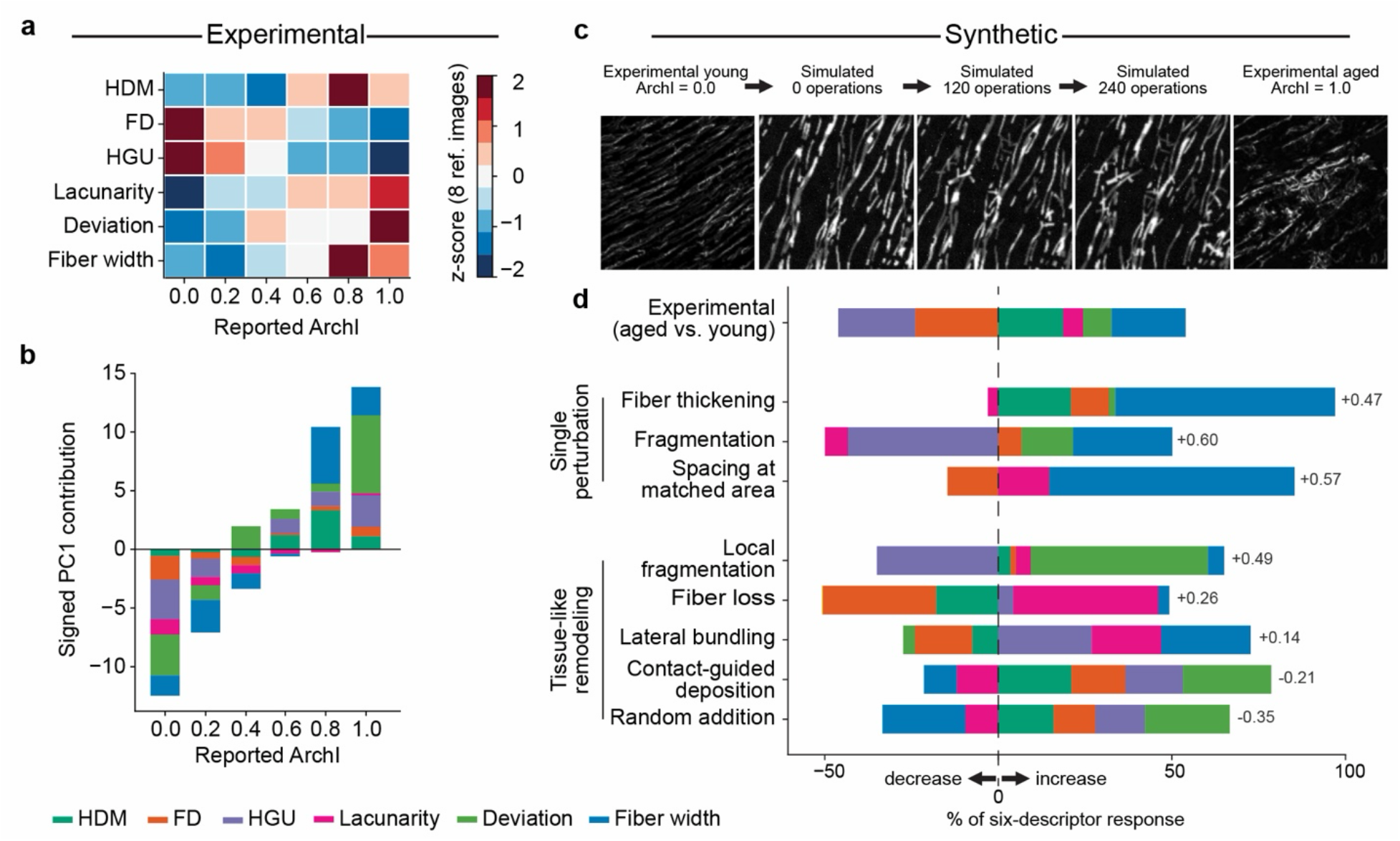
Experimental and synthetic characterization of the Architecture Index (ArchI) across ECM geometries. (**a**) Geometric descriptor profiles corresponding to the experimental images shown in Figure S4, displaying the relative values of HDM, FD, HGU, lacunarity, fiber deviation, and fiber width. (**b**) Decomposition of the corresponding PC1 coordinates into weighted descriptor contributions, showing the contribution of each geometric descriptor to the overall PC1 score underlying ArchI. (**c**) Process-based simulation of ECM remodelling. A tissue-like synthetic network (aligned, wavy, interrupted fibers; initial state calibrated to the experimental tiles) was remodelled by a stochastic mixture of operations (local fragmentation 0.35, lateral bundling 0.25, random rotation 0.15, fiber loss 0.10, thicken / thin 0.15; probabilities of each operation), and rendered 166-pixel crops are shown after 0, 120 and 240 operations (seed 9) between the experimental tiles with the lowest and highest ArchI. Simulated and experimental images were rendered, segmented and measured with the same ridge-based operator. (**d**) Composition of the six-descriptor response for the experimental aged-minus-young direction (heart-level means of standardized descriptors, three hearts per group) and for defined structural changes: single geometric perturbations in controlled geometries (fiber thickening from 4 to 12 px, fragmentation into 8 pieces per fiber, and parallel spacing increased from 32 to 64 px at constant matrix area; 20 paired seeds, truth masks), and remodelling operations applied to the tissue-like network (local fragmentation, fiber loss, lateral bundling, contact-guided deposition, random addition; 30 seeds, 240 operations, tissue-matched operator). Bars give the share of each descriptor in the total scaled response (median paired change divided by the descriptor’s 5th to 95th percentile range); leftward blocks are decreases and rightward blocks increases. The number at the right of each row is the cosine similarity between the row’s response and the experimental direction; leave-one-heart-out ranges are given in Supplementary Figures S3b and S4g.

To examine combinations of structural changes, we simulated a tissue-like network of aligned, wavy, and interrupted fibers whose structural parameters were calibrated to the descriptor profiles of eight experimental image crops (**Supp. Fig. S4a**). Simulated images were rendered as fluorescence-like images and processed alongside the reference crops using the same image analysis procedure within the modeling analysis (**Supp. Fig. S4a,b,d**). Calibration increased the proportion of simulated networks falling within the reference crop range for all six descriptors from 10 % to 39% (**Supp. Fig. S4c**). We then remodeled these networks using fourteen stochastic operations representing structural changes, including lateral bundling, local fragmentation, contact-guided deposition, fiber loss, changes in fiber width, realignment, and coiling (**Supp. Fig. S4e,f**).

A mixture of local fragmentation, lateral bundling, fiber width changes, small rotations, and fiber loss produced partial agreement with the experimental direction, although the response varied across simulations (**Fig. 3c**; median cosine similarity 0.37 across five random seeds; **Supp Fig. S4j,k**). Measured per operation, local fragmentation reduced HGU and raised deviation, fiber loss lowered HDM and FD and raised lacunarity, and lateral bundling raised fiber width, HGU, and lacunarity (**Fig. 3d; Supp. Fig. S4g**). These differing responses illustrate how combinations of structural changes can influence the descriptor profile.

Across 2,000 random process mixtures, 170 met the minimum response-magnitude criterion and showed directional agreement with the experimental pattern (cosine similarity ≥ 0.5; **Supp. Fig. S4h,i)**. These mixtures had a local-fragmentation rate 1.47-fold above the prior median, and rates of random fiber addition and contact-guided deposition 0.44-fold and 0.60-fold of the prior median, respectively. Thickening, bundling, and realignment rates showed no clear enrichment. Within the tested model, these results support local fragmentation as a candidate contributor to the age-associated descriptor pattern. Descriptors measured by Fiji-based measurement routines agreed in rank with the Python implementation for FD, HGU, and lacunarity (Spearman r = 0.86, 0.85 and 0.89, respectively), but not for fiber width (r = -0.01; **Supp. Fig. S4l,m**). The direction of the simulated remodeling response was also not consistently retained across the two measurement pipelines. Synthetic and experimental architectures were therefore compared through descriptor responses without treating their absolute ArchI values as interchangeable.

### Spatial correlative mapping reveals age-associated differences in collagen architecture and tissue stiffness

To relate local ECM architecture to tissue stiffness, we established a correlative mapping workflow that enabled nanoindentation measurements and fluorescence imaging to be spatially mapped within the same tissue regions (**Fig. 4**). A 100-µm thick fresh cardiac tissue slice was mounted on the upper surface of a coverslip containing a patterned polydimethylsiloxane (PDMS) registration grid of labeled 200 µm x 200 µm squares on the opposite surface of the glass (**Fig. 4a**). The underlying grid and its unique coordinate identifiers remained visible through the tissue, providing spatial references for nanoindentation and subsequent relocation of the same myocardial regions during confocal imaging.

**Figure 4.**
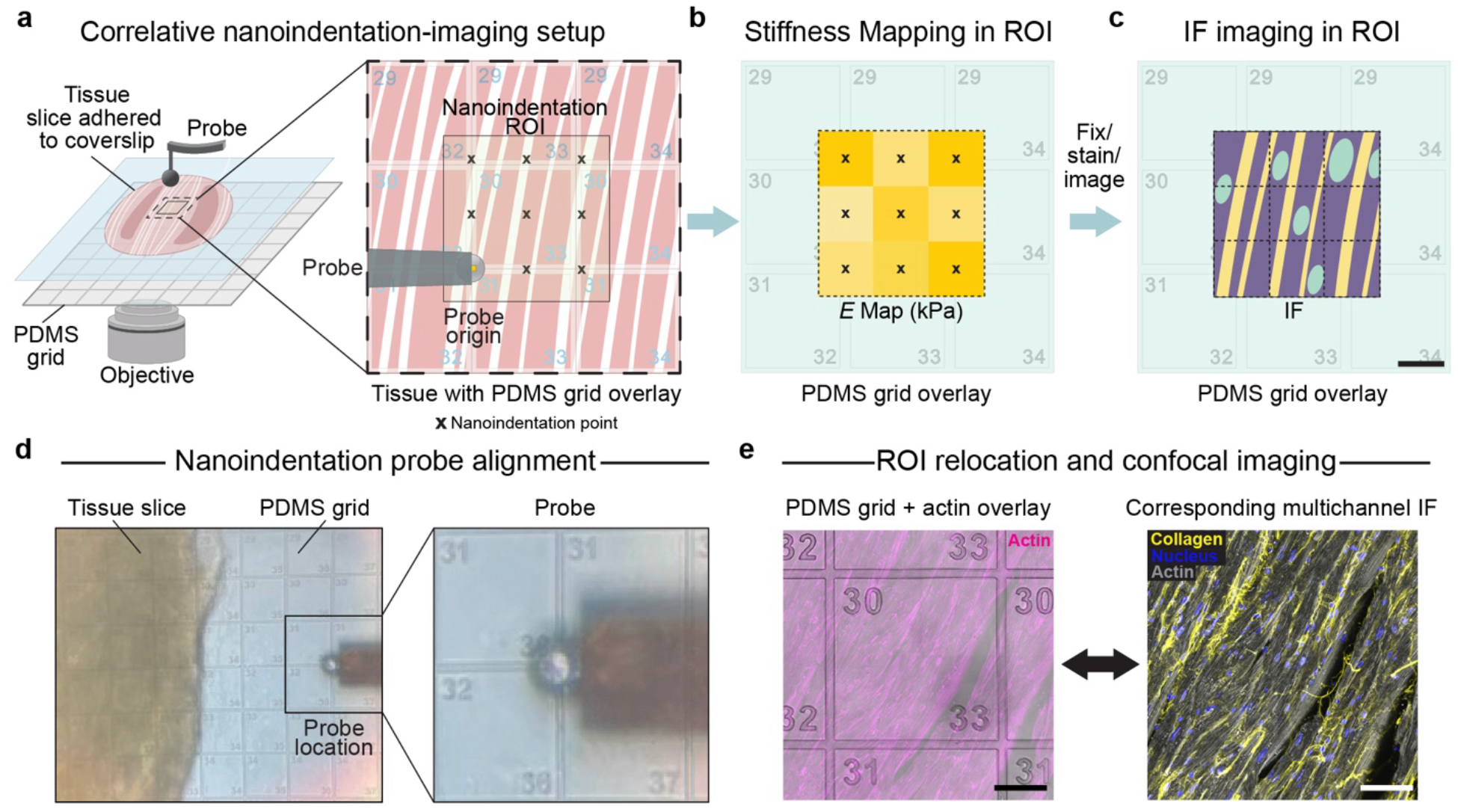
Correlative spatial mapping framework for tissue stiffness and extracellular matrix architecture. (**a**) Schematic illustrating the overall correlative nanoindentation-imaging set-up, with the tissue adhered onto a glass coverslip with a marked PDMS grid underneath. 3x3 nanoindentation ROIs shown with the PDMS grid visible through the tissue slice and probe origin identified in relation to *x-y* coordinate identifiers. (**b**) Stiffness mapping schematic, where nanoindentation measurements were performed within the identified ROI. (**c**) Following indentation, the tissue was fixed and immunofluorescence staining was performed before confocal imaging in the identified ROI. (**d**) Brightfield image of nanoindentation probe alignment with PDMS grid and *x-y* coordinates visible together with the tissue slice (left). Zoom-in of probe location aligned with grid axes (right). (**e**) Matched ROI for confocal imaging, where the same spatial location used for nanoindentation was relocated for confocal imaging. Actin (magenta) overlaid with brightfield of PDMS grid (left) and corresponding multichannel IF image (right) for collagen (yellow), nucleus (blue), and actin (grey). Scale bars: 100 µm (c) and 50 µm in (e).

Local tissue stiffness was first measured in fresh tissue sections using a 3 x 3 nanoindentation array at 100-µm grid point spacing (**Fig. 4b**). Following fixation and immunostaining, the same grid coordinates were used to relocate each nanoindented region for confocal imaging in order to quantify fibrillar collagen ArchI and fluorescence intensity (**Fig. 4c**). The PDMS grid enabled probe alignment during nanoindentation and relocation of the corresponding region for fluorescence imaging **(Fig. 4d,e).**

Consistent with the cryosection findings, aged tissue exhibited significantly higher fibrillar collagen ArchI values than young tissue (**Fig. 5a**; FDR-adjusted *q* = 0.0266), together with increased collagen fluorescence intensity and tissue stiffness in the corresponding regions (**Supplementary Fig. S5b,c**; FDR-adjusted *q* = 0.0266 and 0.0266 respectively; **Supplementary Table S3).** When these measurements were plotted together as registration grid means, young regions predominantly clustered at lower ArchI and stiffness values, whereas aged regions clustered at higher values, with overlap between the groups (**Fig. 5b**). These measurements revealed regional variation within each group alongside the overall age-associated shift in collagen architecture and tissue stiffness.

**Figure 5.**
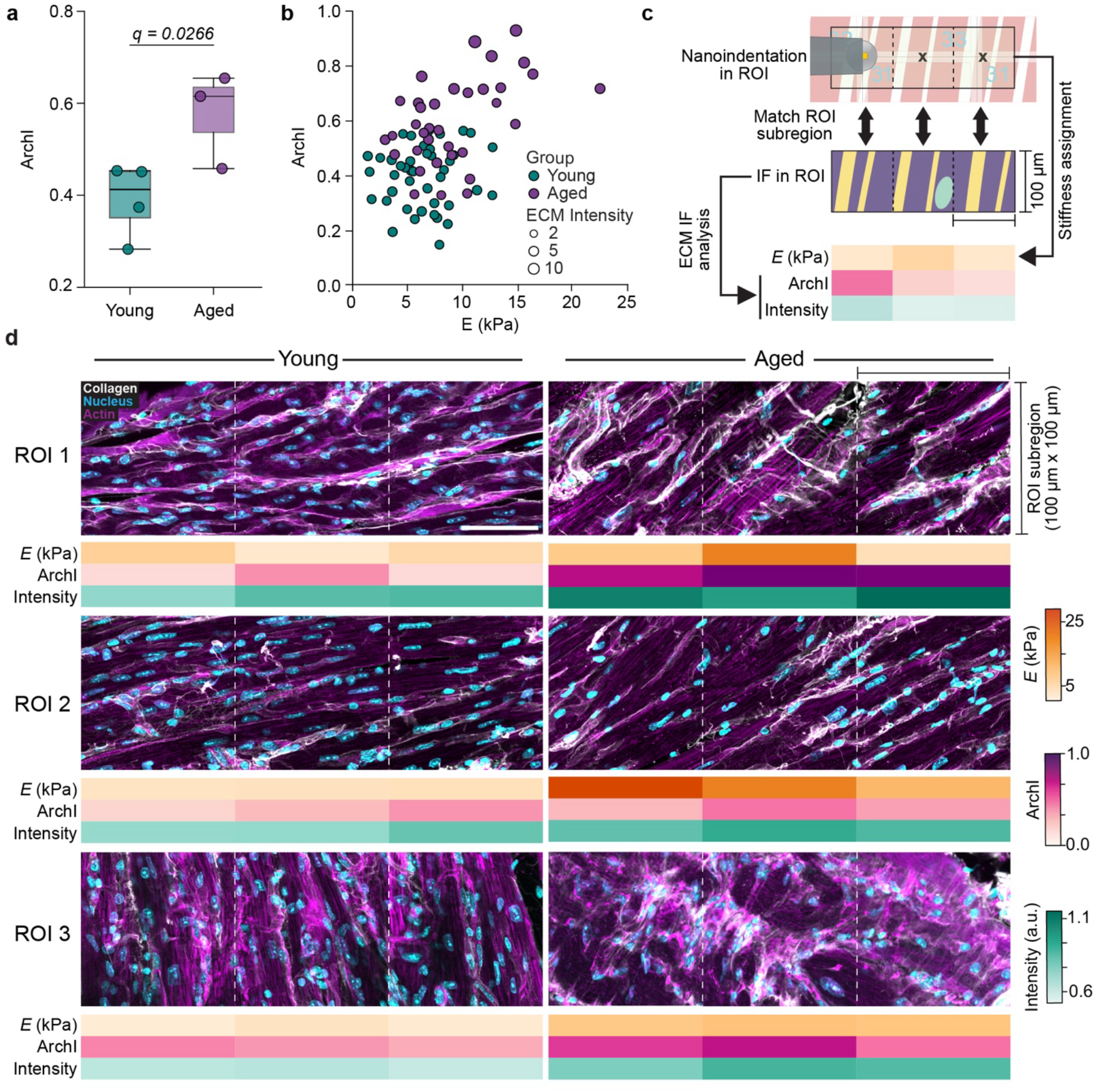
Spatial integration of stiffness, Architecture Index (ArchI), and collagen remodeling in the aging myocardium. (**a**) Boxplots of the ArchI measured in fixed cardiac tissue used for correlative spatial mapping. (**b**) Scatter plot illustrates the relationship between fibrillar collagen ArchI, tissue stiffness (*E*, kPa), and ECM fluorescence intensity. Point size represents nuclei-normalized collagen integrated fluorescence intensity within each matched ROI, and each point corresponds to the mean value of a single spatial registration grid. **(c)** Schematic illustrating analysis within a Grid-Matched ROI, where ECM Analysis was performed on the matched tissue image to obtain ArchI and ECM fluorescence intensity, which were spatially integrated with tissue stiffness measurements from the corresponding 3 × 3 nanoindentation map. Three representative positions from the nine-point map are shown and carried forward for visualization in the subsequent panels. Scale bar, 100 µm. **(d)** Representative grid-matched ROI images (1-3) from young and aged cardiac tissue illustrating spatial heterogeneity in local ECM architecture. Scale bar, 50 µm. Corresponding ECM Analysis shows tissue stiffness (*E*, kPa), ArchI, and ECM fluorescence intensity for the three representative matched positions defined in (c). Group comparisons were performed using ordinary least squares (OLS) regression with cluster-robust (CR1) standard errors. *P* values were adjusted using the Benjamini-Hochberg false discovery rate (FDR) correction, with adjusted *P* values shown in (a). *N* = 4 and 3 biological replicates for young and aged (mice) groups, respectively.

To examine the local heterogeneity underlying these grid-level measurements, we selected three representative regions of interest (ROIs) from each age group, displaying three spatially matched subregions from each nine-point indentation map alongside their corresponding stiffness, ArchI, and collagen fluorescence intensity measurements (**Fig. 5d**). Young regions generally exhibited lower ArchI, fluorescence intensity, and tissue stiffness, with local variations across the selected subregions. In contrast, aged regions exhibited higher values overall, although the spatial distributions of these measurements differed between ROIs. Aged ROI 1 showed elevated ArchI and collagen fluorescence intensity, whereas ROI 2 exhibited the highest stiffness despite comparatively lower ArchI values. ROI 3 displayed elevated ArchI with more moderate stiffness and spatially variable collagen fluorescence intensity (**Fig. 5d**). These observations demonstrate that age-associated increases in collagen architecture and tissue stiffness coexist with local heterogeneity, with the highest stiffness not necessarily corresponding to the highest ArchI or collagen fluorescence intensity.

To further assess the relationship between ECM architecture and tissue stiffness, we performed age-adjusted ordinary least squares regression using the spatially matched measurements. ArchI was positively associated with stiffness after adjustment for age group (β = 6.49 kPa per unit ArchI, 95% CI: 1.46–11.53, *P* = 0.0197; **Supplementary Fig. S5d**). Leave-one-heart-out (LOHO) sensitivity analysis retained a positive ArchI regression coefficient across all seven exclusions, although statistical significance was not maintained in every refit (**Supplementary Fig. S5f; Supplementary Table S11**). An additional regression model incorporating an ArchI × age interaction term revealed a steeper association in aged myocardium (β = 9.44 kPa per unit ArchI, 95% CI: 3.97–14.90, *P* = 0.0055) than in young myocardium (β = 2.19 kPa per unit ArchI, 95% CI: −1.17 to 5.55, *P* = 0.1613), with a significant interaction coefficient (β = 7.25 kPa per unit ArchI, 95% CI: 0.83–13.66, *P* = 0.0326; **Supplementary Fig. S5e**). The interaction coefficient remained positive across all heart exclusions, although its statistical significance was not consistently retained (**Supplementary Fig. S5g–i; Supplementary Table S11).** Overall, these results support a positive association between fibrillar collagen architecture and myocardial stiffness after accounting for age, with a stronger relationship observed in aged myocardium.

## Discussion

Increased tissue stiffness and ECM remodeling are widely recognized features of cardiac aging [11,21,22], yet the extent to which local matrix architecture relates to regional mechanical heterogeneity remains unclear. Here, we derived an integrated Architecture Index (ArchI) from quantitative ECM imaging and combined this analysis with spatial stiffness mapping to examine matrix architecture-mechanics relationships within the same myocardial regions as a function of age. Synthetic geometries further helped interpret how defined structural changes affect the descriptors underlying ArchI. Aged myocardium exhibited distinct remodeling patterns across ECM components and an overall shift toward higher ArchI. Spatially matched measurements also revealed higher fibrillar collagen ArchI and tissue stiffness in aged tissue, with selected local regions displaying different combinations of collagen architecture, fluorescence intensity, and stiffness. These findings highlight component-specific architectural remodeling during cardiac aging and demonstrate an approach for examining its spatial correspondence with tissue mechanics.

Our observations suggest that ECM remodeling during cardiac aging extends beyond a generalized increase in matrix. Individual ECM components exhibited distinct changes in both fluorescence intensity and architecture. Across ECM components, multiple descriptors changed concurrently and often in different directions, indicating that measurements of matrix occupancy, fiber geometry, or network continuity each capture only part of the remodeling pattern. Consistent with this, the component-specific PCA showed that age-associated architectural differences involved different combinations of descriptors (**Fig. 1c**), while pairwise correlations showed that relationships among these features also varied across ECM components (**Fig. S1c**). Integrating these descriptors into ArchI provides a summary of the dominant architectural variation within a shared score. The underlying descriptors remain necessary to identify which structural features contribute to the architectural differences observed in each ECM component.

By representing each ECM component using the same set of structural descriptors, interstitial, pericellular, and basement membrane-associated ECM networks can be compared. In experimental examples, higher ArchI scores were generally accompanied by lower HGU and greater apparent fiber width. However, similar scores can arise from different combinations of descriptors. ArchI therefore summarizes architectural variation within the reference dataset, determined by features, weights, and scaling. This integrative approach has parallels in established histological scoring systems [23] and quantitative approaches applied to pulmonary fibrosis [24], liver disease and fibrosis [25–27], and myocardial fibrosis [28]. Independent testing using fixed descriptor weights and scaling will be needed to determine how well ArchI generalizes beyond the present samples and imaging conditions.

The simulations helped interpret how defined structural changes affect the descriptors underlying ArchI. Controlled geometries showed distinct responses to thickening, fragmentation, and spacing changes, each reproducing only part of the aged-minus-young difference. On a tissue-like network calibrated to the experimental image crops, mixtures of remodeling operations produced partial and variable agreement with the experimental descriptor pattern. Local fragmentation was enriched among mixtures meeting the response magnitude and directional agreement criteria, supporting it as a contributor within the tested model. These results suggest that changes in network continuity can contribute to the observed architectural pattern, although the biological processes responsible remain to be established experimentally [29]. The Python-based measurements used in the synthetic analysis showed good rank agreement with the corresponding Fiji measurements for FD, HGU, and lacunarity, but not for fiber width. Furthermore, the direction of simulated remodeling was not consistently retained across the two measurement pipelines. The simulations therefore provide a controlled means of examining descriptor sensitivity to structural changes, while these measurement differences highlight the limitations of applying the model to experimental ArchI.

In tissue, interpreting ArchI alongside the individual structural descriptors and fluorescence intensity measurements helps distinguish how different ECM components remodel with age. Fibrillar collagen showed higher ArchI and a trend toward increased fluorescence intensity, consistent with concurrent changes in matrix content and organization [6,30]. Fibronectin similarly exhibited architectural remodeling despite no significant difference in fluorescence intensity, consistent with its mechanosensitive assembly and central role in collagen fibrillogenesis and matrix organization [31–34]. Among basement membrane components, laminin fluorescence intensity decreased while ArchI increased, whereas collagen IV showed increased fluorescence intensity without a significant ArchI difference. These contrasting responses occurred within matrix networks that jointly contribute to basement membrane organization [35–37]. Lastly, collagen VI showed reduced fiber orientation dispersion without a significant change in ArchI, illustrating how individual geometric features can change without a detectable shift in the composite score [38–41].

Correlative approaches such as STIFMap have combined spatially registered atomic force microscopy measurements with collagen and nuclear imaging to predict stromal stiffness in breast tumors [20]. Although representative aged regions of our samples exhibited distinct combinations of these measurements, age-adjusted regression revealed a positive association between ArchI and apparent Young’s modulus. The ArchI × age interaction further indicated a steeper association in aged myocardium than in young myocardium, suggesting that the relationship between ECM architecture and tissue stiffness may differ with age. The interaction coefficient remained positive across leave-one-heart-out analyses, although statistical significance was not consistently retained. These findings highlight the importance of spatially resolved measurements in identifying architecture-stiffness relationships that may be obscured by regional heterogeneity. However, given the limited number of independent hearts and the observational nature of the analysis, we cannot conclude that collagen architectural remodeling alone caused the observed tissue stiffening.

Local architectural differences may also influence the matrix cues encountered by resident cardiac cells [18,41–43]. For example, aligned collagen-mimetic topographies were shown to promote cardiac myofibroblast differentiation through p38-YAP-TEAD signaling [44]. In dermal fibroblasts, three-dimensional fibrin matrices with independently tunable architecture and bulk stiffness showed that network architecture influences cell survival and differentiation [45]. Engineered myocardial microtissues further demonstrate that matrix fiber alignment and stiffness affect cardiomyocyte organization and contractility [16]. Lastly, our DECIPHER hybrid biomaterial showed that young ECM ligand presentation can promote cardiac fibroblast quiescence despite an aged-like stiffness cue [6,7]. These findings motivate investigating whether the architectural and mechanical differences observed here generate distinct local cellular responses during cardiac aging.

However, several limitations should be considered. The correlative design identifies spatial associations between ECM architecture and tissue stiffness but does not establish whether specific matrix features directly cause local stiffening. Perturbation studies targeting specific ECM components, matrix remodeling pathways, or stiffness-modifying mechanisms will be required to test causality. The imaging-based architectural measurements were derived from tissue sections and may not fully represent three-dimensional ECM organization. Further, the analysis focused on selected ECM components and image-derived descriptors, and therefore does not capture the full myocardial matrisome, non-matrix contributors to tissue mechanics, or biochemical matrix modifications such as collagen crosslinking, glycation, and other post-translational changes. Although ArchI provides a data-driven integrated readout, it depends on the selected structural features and reference dataset used for scaling and weighting and should therefore be interpreted alongside its underlying descriptors. Lastly, the synthetic model calibration limits the interpretation of physical fiber dimensions and the range of architecture represented by the model.

Together, these findings establish a quantitative description of component-specific ECM remodeling and a means to examine collagen architecture alongside local tissue mechanics during cardiac aging. More broadly, the measured architectural profiles and corresponding stiffnesses may guide the development of biomaterial platforms that reproduce selected features of young and aged myocardial environments [46]. Combining these platforms with targeted perturbations and cellular readouts could help determine which matrix changes contribute to altered cardiac cell behavior and tissue dysfunction.

## Acknowledgements

The authors thank Jennifer Marlena (MBI, NUS) for illustrations; Low Boon Chuan, Yan Jie, Hepi Heri Susapto, Hashan Preet Singh (MBI, NUS), Matthew Ackers-Johnson, Roger Foo (CVRI, NUS), Cesare Mellace, and Lara Pereira (Optics11) for fruitful discussions and technical advice; Chii Jou Chan, Ng Boon Heng, Leong Kim Whye, and Arikta Biswas (MBI, NUS) for assistance with animal handling and vibratome usage; and Chii Jou Chan and Rong Li (MBI, NUS) for providing mouse heart tissues via the NUS Tissue Sharing Program. We thank the MBI Wet Lab Core for facility and technical support, and the Singapore Microscopy and Bioimaging Analysis (SiMBA, MBI) Core for microscopy and data processing facilities.

## Author contributions

J.L.Y. and M.F.H.R. conceptualized the project and wrote the manuscript. J.L.Y. supervised data collection and analysis. M.F.H.R. performed the majority of experiments and data analysis. A.G. developed the synthetic index model. R.H., A.R.S., X.S., K.N. and G.G. assisted with bench experiments. A.G., H.T.O., Y.L., O.H.T., M.S., T.H. assisted with data analysis.

## Code availability

The Python code used for nanoindentation analysis and Hertz model fitting is publicly available on GitHub at https://github.com/jennyounglab/NI-MPD_CardiacMechanics. The Fiji/ImageJ macros and Python scripts used in this study are available at https://github.com/jennyounglab/ArchI-Stiffness. These include codes for ECM architectural descriptor quantification, Architecture Index (ArchI) derivation, spatial matching of ECM architecture with local tissue stiffness measurements, and statistical analysis of architecture–stiffness relationships. The repository also contains code for synthetic ECM analyses, including controlled geometric perturbations, tissue-like stochastic network generation, model calibration, simulated ECM remodeling, and ArchI validation. The ArchI-Stiffness repository is currently private during manuscript review and will be made publicly accessible upon publication. Access to the code during peer review can be arranged through the corresponding author.

## Data Availability

The datasets supporting the findings of this study, including ECM architectural descriptors, Architecture Index (ArchI) measurements, fluorescence intensity measurements, and spatially matched tissue stiffness data, are available at https://github.com/jennyounglab/ArchI-Stiffness.

## Materials and Methods

### Tissue isolation and preparation

Hearts from young (25 days to 1 month) and aged (18–24 months) female C57BL/6J mice were obtained through the NUS Animal Tissue Sharing Programme. Following euthanasia by carbon dioxide (CO₂) inhalation, hearts were immediately immersed in sterile Tyrode’s solution supplemented with 30 mM 2,3-butanedione monoxime (BDM; Merck, B0753-100G) to minimize contraction and preserve tissue integrity. Excess connective and major vascular tissues were carefully removed using sterile surgical instruments. A 4% (w/v) low–melting point agarose solution (Invitrogen, 16520) prepared in sterile 1× PBS was maintained at 37 °C prior to embedding. Hearts were embedded in agarose and oriented along the sagittal plane to obtain longitudinal sections, then placed on ice to allow solidification [47]. All agarose-embedded hearts prepared for subsequent cryosectioning were sectioned at a thickness of 1 mm using a Leica VT1200S vibratome at a cutting speed of 0.16 mm s⁻¹.

### Cryo-embedding and sectioning

The 1 mm-thick cardiac tissue slices were fixed in 4% paraformaldehyde (PFA) for 1h at room temperature, followed by washing with 1× phosphate-buffered saline (PBS) to remove residual PFA. The slices were subsequently embedded in optimal cutting temperature (OCT) compound, rapidly frozen in liquid nitrogen, and stored at −80 °C until sectioning. OCT-embedded tissues were sectioned at a thickness of 10 µm using a cryostat (Leica CM1950). Sections were collected onto Superfrost™ Plus microscope slides (VWR) and stored at −80 °C until immunofluorescence staining.

### IHC and imaging

Prior to staining, cryosections were rehydrated in 1× PBS to remove residual OCT. Samples were then permeabilized and blocked 30 min at room temperature in 2% bovine serum albumin (BSA; Sigma-Aldrich) containing 0.1% Triton X-100. Primary antibodies against the selected ECM components were incubated overnight at 4 °C in blocking solution at the dilutions listed in **Supplementary Table S8**. Following three washes in 1× PBS, sections were incubated with the appropriate secondary antibodies, CF® phalloidin (Biotium) for F-actin staining, and the CNA35 collagen-binding probe (1 µM) for 1 h at room temperature in the dark. The EGFP–CNA35 fusion protein was expressed from the pET28a-EGFP-CNA35 plasmid (Addgene #61603) and purified before use. Nuclei were counterstained with Hoechst 33342 (Invitrogen), and samples were mounted using Fluoromount-G™ (Invitrogen).

Samples were imaged using an Olympus FV3000 laser-scanning confocal microscope with a 30× silicon immersion objective and 2× digital zoom. Low-magnification overview images were first acquired using a 4× objective to map the myocardium in longitudinally oriented cardiac tissue sections and identify the left ventricular wall, right ventricular wall, and interventricular septum. Approximately 10 spatially distributed, nonoverlapping fields of view (212 × 212 µm) were selected within the myocardium of each cardiac region, using the overview images as reference maps to guide sampling locations and avoid repeated imaging of the same tissue areas. Z-stacks consisting of six optical sections acquired at 1 µm intervals, spanning a total depth of 5 µm, were merged into two-dimensional images for subsequent analysis. Images containing technical artifacts or insufficient image quality were excluded from subsequent analyses. For each staining condition, laser power, detector settings, and other intensity-related acquisition parameters were kept constant across samples to enable quantitative comparisons of fluorescence intensity between experimental groups. Images were subsequently imported into Fiji/ImageJ (version 1.54p) for quantitative image analysis.

### ECM architecture quantification

Extracellular matrix (ECM) architecture was quantified from confocal fluorescence images using TWOMBLI (The Workflow Of Matrix BioLogy Informatics), implemented in Fiji/ImageJ (version 1.54p) [19]. Images were processed using the TWOMBLI workflow, including contrast adjustment, ridge detection, and generation of ECM network masks, with the analysis parameters provided in Supplementary Table S9. Four architectural descriptors were extracted using TWOMBLI: high-density matrix (HDM), fractal dimension (FD), lacunarity, and hyphal growth unit (HGU). HDM was quantified from intensity-thresholded HDM masks, whereas FD, lacunarity, and HGU were derived from ridge-detection masks. HGU was calculated as skeleton length per endpoint and used as a proxy for network continuity. Fiber deviation and fiber width were quantified separately from segmentation masks using a custom Fiji/ImageJ macro, with representative segmentation masks shown in **Supplementary Fig. S1B**. Fiber deviation was calculated as the major axis-weighted mean angular deviation of segmented ECM structures from their predominant axial orientation, with angular differences constrained to 0–90° and expressed in degrees. Apparent fiber width was calculated as the major axis-weighted mean minor-axis length of these structures and expressed in micrometers (µm). Detailed definitions and interpretations of each architectural metric are provided in **Supplementary Table 6**.

ECM fluorescence intensity was quantified from 8-bit grayscale fluorescence images using Fiji/ImageJ. Background subtraction was performed with sliding paraboloid option. ECM-positive regions were subsequently identified using the Triangle automatic thresholding method, followed by particle analysis to quantify raw integrated density (RawIntDen) and segmented area. For the cryosection dataset, area-normalized fluorescence intensity was calculated by dividing the total RawIntDen of ECM-positive regions by their corresponding total area (µm²).

### Individual ECM component PCA and correlation analysis

To examine component-specific patterns of ECM architectural variation, principal component analysis (PCA) was performed separately for fibrillar collagen, fibronectin, laminin, collagen IV, and collagen VI using the six architectural descriptors. For each ECM component, image-level measurements were averaged within individual hearts to obtain one mean value per descriptor, with each heart representing an independent biological replicate. The resulting heart-level measurements from Young and Aged hearts were pooled and standardized using Z-score normalization before PCA to account for differences in numerical scales. The first two principal components (PC1 and PC2) were used to visualize architectural variation between age groups. Pairwise correlations among the six architectural descriptors were calculated using Pearson’s correlation coefficients based on pooled Young and Aged heart-level mean measurements for each ECM component.

### Architecture Index (ArchI) derivation

To summarize the dominant variation in ECM architecture, an Architecture Index (ArchI) was computed from six structural descriptors: high-density matrix (HDM), fractal dimension (FD), lacunarity, hyphal growth unit (HGU), fiber deviation, and fiber width. Each metric was standardized by Z-score normalization within each ECM component using the mean and standard deviation calculated from pooled Young and Aged image-level measurements for that component, to ensure comparability across features with different scales.

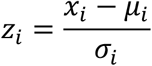

where *x_i_* is the original value of the *i*-th architectural metric, *μ_i_* and *σ_i_* are the corresponding mean and standard deviation, respectively, calculated from young and aged image-level measurements within each ECM component.

Following normalization, the standardized measurements were pooled across ECM components and aggregated at the heart level. For each heart, the median value of each architectural descriptor was calculated across all images to reduce the influence of extreme values and localized variability. A second Z-score normalization was then applied to the resulting heart-level feature matrix using the mean and standard deviation of each descriptor calculated across all Young and Aged hearts. This ensured that differences in descriptor variance did not disproportionately influence the subsequent principal component analysis (PCA).

PCA was then performed on the standardized heart-level feature matrix. The loadings of the first principal component (PC1), which captured the dominant axis of architectural variation (**Supp. Table 4 &5**), were used as weighting coefficients to calculate the PC1-derived Architecture Index:

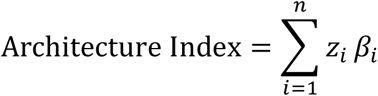

where *z_i_* represents the standardized (z-score) value of the *i*-th architectural metric, *β_i_* denotes the corresponding loading of the *i*-th metric on the first principal component (PC1), and *n* is the number of architectural metrics included in the analysis.

This formulation integrates multiple structural features into a single variable representing the global architectural state of the ECM. For consistent biological interpretation, the index was oriented such that higher values correspond to aging-associated ECM remodeling by enforcing a consistent sign direction based on group-level PC1 trends. The index was subsequently normalized to a bounded range using min-max scaling. This reference-based scaling was subsequently applied to the correlative stiffness-structure datasets, enabling direct comparison of Architecture Index values within a consistent quantitative framework. Under this convention, higher values indicate a more aged-like ECM phenotype.

### Preparation of the correlative mapping workflow

Glass coverslips (15 × 15 mm) were cleaned by sequential sonication in acetone, ethanol, and Milli-Q water for 10 min each, air-dried, and treated with ultraviolet/ozone for 5 min. Polydimethylsiloxane (PDMS; Sylgard 184, Dow) was prepared by mixing the base and curing agent at a 10:1 (w/w) ratio. The PDMS mixture was poured onto a custom microfabricated grid wafer and spin-coated at 1000 rpm for 30 s to produce a uniform thin PDMS layer. A cleaned glass coverslip was then carefully placed onto the uncured PDMS, followed by vacuum degassing to remove trapped air bubbles. The assembly was cured at 80 °C for 1.5 h, after which the bonded PDMS grid and glass coverslip were removed from the wafer. The resulting platform contained a labeled PDMS registration grid on one face of the coverslip, with the opposite glass face available for tissue mounting.

### Tissue preparation

Fresh, agarose-embedded hearts were sectioned at a thickness of 100 µm using a Leica VT1200S vibratome at a cutting speed of 0.10 mm s⁻¹. Following sectioning, the surrounding agarose was carefully removed, and excess buffer was wicked from the edges of each tissue slice using a Kimwipe to facilitate mounting. A thin layer of tissue adhesive (Vetbond™ Tissue Adhesive, 1469SB) was applied to the glass surface opposite the PDMS registration grid. Tissue slices were mounted onto the adhesive-coated surface and allowed to adhere for approximately 5 min, after which the platform was submerged in Tyrode’s solution supplemented with 30 mM 2,3-butanedione monoxime (BDM), as described above. Slices were maintained in an ice bath, and mechanical characterization was initiated within 2 h of sectioning.

### Nanoindentation

Fresh cardiac tissue slices were mechanically characterized using an Optics11 Life Chiaro Nanoindenter (Optics11 Life, The Netherlands) equipped with a spherical probe (spring constant, 0.5 N m⁻¹; tip radius, 50 µm). To minimize probe–sample adhesion, the probe was coated with 1% (w/v) Pluronic F-127 solution (Sigma-Aldrich, P2443) before each experiment. Probe calibration was performed against a glass substrate before establishing the measurement origin on the PDMS registration grid using the grid boundaries as spatial reference markers. Indentation maps were acquired over predefined regions of interest (ROIs) using a 3 × 3 indentation array with a center-to-center spacing of 100 µm and an indentation depth of 3 µm. Measurements were performed in Tyrode’s solution supplemented with 30 mM 2,3-butanedione monoxime (BDM) at room temperature, with loading and unloading rates of 1 µm s⁻¹ and a dwell time of 1 s at maximum indentation depth. Each ROI was assigned a unique grid coordinate (x, y), which was retained for subsequent spatial registration with fluorescence imaging.

Young’s modulus was calculated from force–indentation curves using a custom Python analysis pipeline, adapted from previously published methods [48], based on Hertzian contact mechanics with an assumed Poisson’s ratio of 0.5 [49]. Force-indentation curves were subjected to baseline correction and Savitzky–Golay smoothing before fitting. The loading curve was fitted over an indentation range of 0.5–2.5 µm, with the initial contact region identified using a force threshold based on baseline fluctuations and the contact-point offset subsequently optimized during Hertz model fitting. The precision of the fitted Hertz coefficient was evaluated using the relative standard error, defined as the square root of the corresponding covariance matrix diagonal element divided by the fitted coefficient

### Correlative stiffness-architecture imaging

Following nanoindentation, the 100-µm tissue slices were fixed in 4% paraformaldehyde (PFA) for 15 min at room temperature and immunostained as described above. Confocal imaging was performed using an Olympus FV3000 laser-scanning confocal microscope under the same acquisition parameters described for fluorescence imaging. Transmitted-light brightfield imaging was used to identify the PDMS registration grid and relocate the previously indented regions of interest (ROIs) using the recorded grid coordinates (x, y). A 2 × 2 confocal image mosaic was acquired for each ROI with 10% overlap between adjacent fields and stitched using the Olympus FV31S-SW software. The stitched images were subsequently processed using a custom ImageJ macro to straighten the grid boundaries, crop a central region measuring 300 × 300 µm centered on the PDMS registration grid, and subdivide the cropped region into a 3 × 3 array of 100 × 100 µm subregions corresponding to the individual indentation positions.

Following image acquisition, each fluorescence image was matched to its corresponding nanoindentation dataset based on the recorded grid coordinates. Within the spatially matched regions, apparent Young’s modulus (E, kPa) obtained from nanoindentation was paired with ArchI and nuclei-normalized collagen integrated fluorescence intensity calculated from the corresponding image subregions. Collagen integrated fluorescence intensity (RawIntDen) was normalized by the number of nuclei within each subregion (RawIntDen/Nuc). These measurements were integrated into a single correlative dataset for subsequent spatial analyses. Grid-level means were used for joint visualization in Fig. 5b, with each 3 × 3 grid representing one measurement region within an individual heart. For Fig. 5d, three selected positions from each representative nine-point indentation grid were displayed to illustrate local spatial variations in ECM architecture and tissue stiffness. These representative grids were selected from a subset of biological replicates rather than all hearts analyzed.

### Simulated ECM architectures

Controlled geometries: Two-dimensional ECM masks were generated in paired controlled experiments in which one structural parameter was varied at a time while the others were held constant (fiber width, parallel spacing with and without matched matrix area, gap heterogeneity, angular dispersion, crossing angle, fragmentation, line dropout, branching depth and lattice spacing). Each level used the same latent random seed as its baseline, so that responses are paired differences; 20 seeds were used per experiment on a 256 × 256 pixel field, and finite segments were rasterized as supersampled oriented rectangles thresholded at 50% coverage. Generation acceptance required less than 1% latent clipping, prespecified topology for fixed-topology experiments and at least 80% unique masks per level.

Tissue-like process model: Tissue-like networks were generated as sets of wavy lamellae with a von Mises orientation distribution [50], Gamma-distributed spacing, log-normal widths, a sinusoidal plus random-walk centreline [51], Poisson-distributed interruptions, cross-bridges and short aligned fragments, on a 384 × 384 pixel field. The eleven hyperparameters of this initial state were calibrated to the eight experimental reference tiles by rejection approximate Bayesian computation [52]: 3,000 networks were drawn from uniform prior ranges, rendered and measured, and the 60 draws closest to the median tile descriptor vector in scaled six-descriptor space were accepted; the posterior medians defined the calibrated start (Supp. Figure S4a). Networks were remodeled by a stochastic process in which fourteen operations were applied in random order to uniformly chosen fibers with probabilities proportional to a rate vector[53,54] : eight geometric operations (rotate, translate, thicken or thin, break, bend, remove, add, branch) and six biological operations that use the same primitives with parameters computed from the current network (realignment towards the local director[29], coiling, lateral bundling of neighbouring fibers, cross-bridging, contact-guided deposition of a new fiber next to a template fiber [55,56], and local fragmentation of a segment into short disordered pieces). Every operation was recorded so that any network can be regenerated exactly. Per-operation fingerprints were obtained by applying one operation type for 240 steps to 30 calibrated networks and expressing the change of each descriptor per 100 operations.

Rendering and measurement: Latent networks were rasterized as supersampled polygons and rendered as fluorescence-like images with a distance-transform intensity profile, short-scale labelling variation, speckle and Gaussian noise. Rendered images and experimental reference tiles were segmented with the same operator: percentile normalization, a multi-scale Sato tubeness filter [57] over scales of 1 to 2 pixels, Otsu thresholding of the positive response [58] and removal of objects below 12 pixels; descriptors were computed on 166-pixel crops matching the tile size. The six descriptors were foreground area fraction (HDM), box-counting fractal dimension (FD), skeleton length per endpoint (HGU)[59], gliding-box lacunarity, structure-tensor orientation deviation [60] and area-over-length fiber width; they correspond in definition to the TWOMBLI descriptors [19,61] but were kept in their own units. For the cross-check with the experimental Fiji/ImageJ measurement routines, AnaMorf (headless Fiji) provided FD, HGU and lacunarity on the same ridge masks, and ImageJ particle analysis reproduced the experimental deviation-and-width workflow on the same images.

Comparison with experimental ageing: Synthetic responses were expressed as paired changes from the initial network divided by each descriptor’s 5th to 95th percentile range across the study, and their composition was displayed as the share of each descriptor in the total absolute response. The experimental reference was the aged-minus-young difference between heart-level means of standardized descriptors (three hearts per group; the standardization and PC1 loadings of Supp. Table 4). Directional agreement was the cosine similarity between a synthetic response and this direction, with leave-one-heart-out recalculations as sensitivity ranges. Synthetic values were never passed through the experimental scaler, and synthetic and experimental ArchI values are not compared directly. To identify processes consistent with the experimental direction, 2,000 rate vectors were drawn log-uniformly over two decades for all fourteen operations, simulated for 240 operations from the calibrated start, and the median log-rate of each operation among mixtures with cosine of at least 0.5 (and, separately, at most −0.5) was compared with the prior median, with 5th to 95th percentile bootstrap intervals. Rejection ABC with local-linear regression adjustment[52] on the mean descriptor vectors of the four lowest- and four highest-index tiles gave posterior rate distributions for young-like and aged-like structure. Realism of the synthetic networks was quantified against the tiles by the energy distance [62], the median nearest-neighbour distance, support and precision in scaled descriptor space. Simulations were implemented in Python (numpy, scipy, scikit-image, OpenCV); code and configurations are available at https://github.com/jennyounglab/ArchI-Stiffness/tree/main/ArchI_Modelling.

### Statistical analysis

Statistical analyses and data visualization were performed using custom Python scripts (Python 3.9.7) with NumPy (version 2.0.2), pandas (version 2.3.2), SciPy (version 1.13.1), statsmodels (version 0.14.5), scikit-learn (version 1.6.1), Matplotlib (version 3.9.4), seaborn (version 0.13.2), and Plotly (version 6.3.0). The openpyxl package (version 3.1.5) was used for spreadsheet input and output. Individual mice constituted independent biological replicates, with fluorescence images and registration grids representing repeated measurements nested within each heart. Data are presented as mean ± standard deviation (SD) for individual measurements or as group means with heart-level 95% confidence intervals (CIs), where indicated (Figs. 1, 2, and 5; Supplementary Fig. S5; Supplementary Tables 1–3 and 10). The level of aggregation and definitions of graphical summaries are specified in the corresponding figure legends and supplementary table captions. The numbers of independent hearts and within-heart measurements for each analysis are provided in Supplementary Table 12.

Data are presented as mean ± standard deviation (SD) or as group means with heart-level 95% confidence intervals (CIs), where indicated. Heart-level 95% CIs were calculated from the means of independent biological replicates, whereas statistical comparisons were performed using image-level ordinary least squares (OLS) regression with mouse-clustered (CR1) standard errors, as described below.

Comparisons between young and aged groups in Figs. 1 and 2 were performed using OLS regression with age group as the predictor and individual fluorescence images as the observational units. Image-derived ECM fluorescence intensity, architectural descriptors, and Architecture Index (ArchI) values were analyzed as the respective outcome variables. Cluster-robust (CR1) standard errors were estimated with biological replicate (mouse) specified as the clustering variable to account for repeated measurements obtained from the same heart. Statistical inference was based on a two-sided Student’s t-distribution with degrees of freedom equal to the number of independent clusters minus one (df = 5 for six hearts). Regression coefficients (β) represent the estimated mean difference between aged and young groups (Aged − Young). Exact two-sided *P* values are reported, and statistical significance was defined as *P* < 0.05.

For the correlative mapping analyses, spatially matched ArchI, collagen fluorescence intensity, and tissue stiffness measurements were averaged within each registration grid. Group differences between young and aged tissue were evaluated using OLS regression with age group as the predictor, using the same mouse-level clustering and inferential procedure described above. To account for multiple grid measurements obtained from the same biological replicate, cluster-robust (CR1) standard errors were estimated with biological replicate specified as the clustering variable while retaining all grid-level observations in the model. Regression coefficients (β) represent the estimated mean difference between aged and young groups, and exact two-sided *P* values were calculated from the cluster-robust model. Per-heart means were additionally computed as descriptive summaries of biological replicate-level central tendencies. To account for multiple comparisons across the Architecture Index, fluorescence intensity, and tissue stiffness, *P* values for the three correlative outcomes were adjusted using the Benjamini-Hochberg false discovery rate (FDR) procedure. For comparisons across ECM components and individual architectural descriptors, *P* values were adjusted for multiple comparisons using the Benjamini–Hochberg false discovery rate (FDR) procedure, with adjusted *P* values (*q*) < 0.05 considered statistically significant.

To assess the association between ECM architecture and tissue stiffness, spatially matched measurements were analyzed using OLS regression with ArchI as a continuous predictor and age group as a categorical covariate. An additional model incorporating an ArchI × age interaction term was fitted to determine whether the ArchI–stiffness association differed between young and aged myocardium. Both models used mouse-clustered CR1 standard errors, with two-sided statistical inference based on a Student’s t-distribution with degrees of freedom equal to the number of independent hearts minus one (df = 6 for seven hearts). For the interaction model, age-specific regression slopes were estimated, with the aged slope and its 95% CI calculated using the full cluster-robust covariance matrix. Model robustness was assessed using leave-one-heart-out (LOHO) sensitivity analyses, in which all measurements from one heart were sequentially excluded and the models refitted. Spearman’s rank correlation analysis was performed separately for young and aged myocardium to assess associations between ArchI and tissue stiffness. Regression coefficients, 95% CIs, and exact two-sided *P* values were reported for the full and reduced models, while Spearman’s correlation coefficients (ρ) and corresponding two-sided *P* values were reported separately without adjustment for within-heart clustering (Supplementary Fig. S5; Supplementary Table 11).

### AI-assisted code development

GitHub Copilot (GitHub) was used to assist with code development and debugging. All analytical procedures, computational parameters, and generated outputs were reviewed and validated by the authors.

## Supplementary Figures

**Figure S1.**
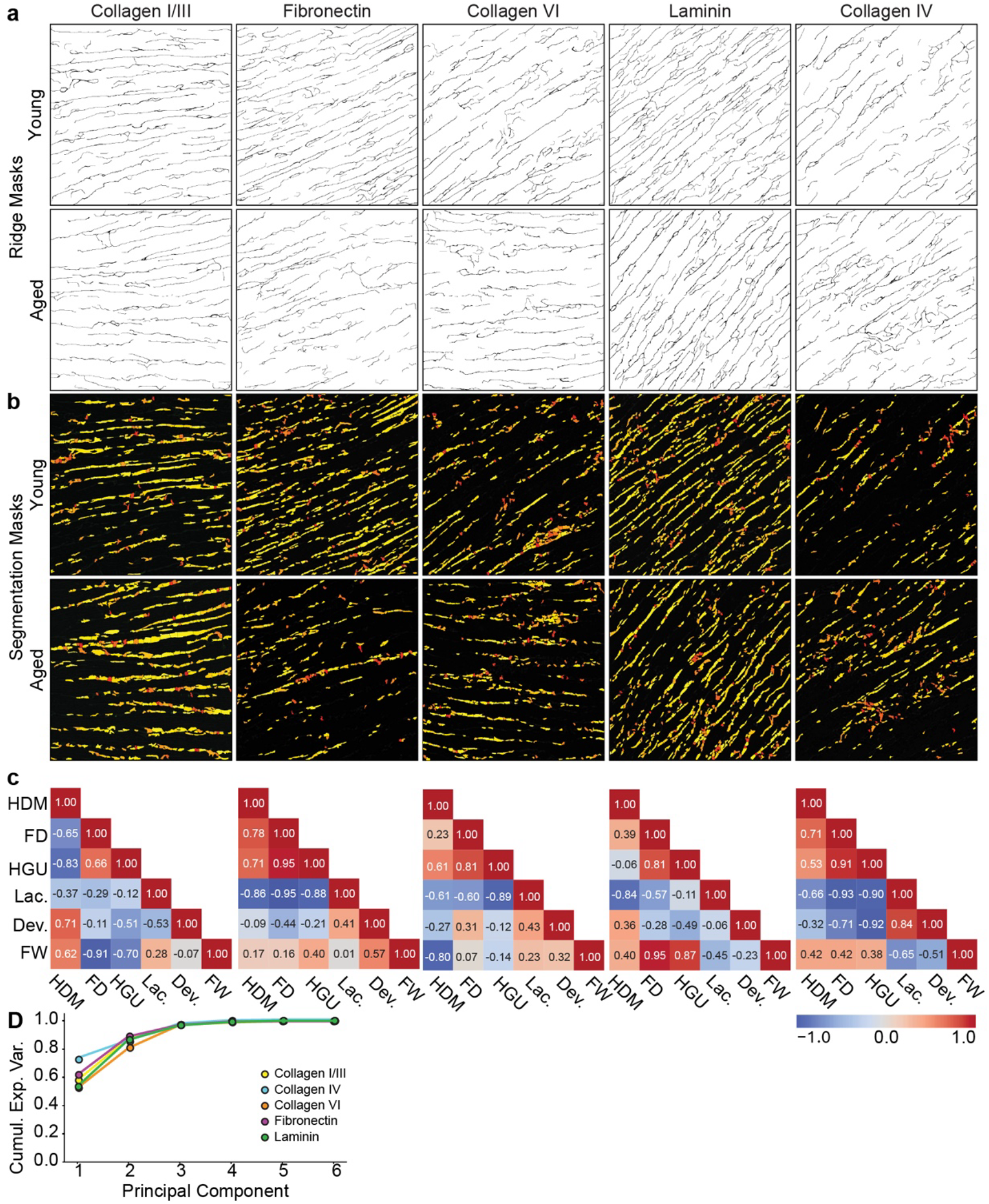
Image processing and PCA validation for quantitative ECM architectural analysis. Representative image-processing outputs used for the quantification of ECM architectural parameters across all ECM components. **(a)** Ridge masks generated using the published TWOMBLI Fiji macro were used to quantify fractal dimension (FD), hyphal growth unit (HGU), and lacunarity, whereas high-density matrix (HDM) was quantified separately using TWOMBLI-generated HDM masks. **(b)** Segmentation masks generated using a custom Fiji/ImageJ pipeline were used to quantify fiber width (FW) and deviation (Dev.). **(c)** Correlation heatmaps of ECM architectural features for individual ECM components in young and aged murine cardiac tissue. Correlation coefficients are displayed within each matrix. *n* = 3 biological replicates (mice) per group. **(d)** Cumulative explained variance of the principal component analysis (PCA) for each ECM component.

**Figure S2.**
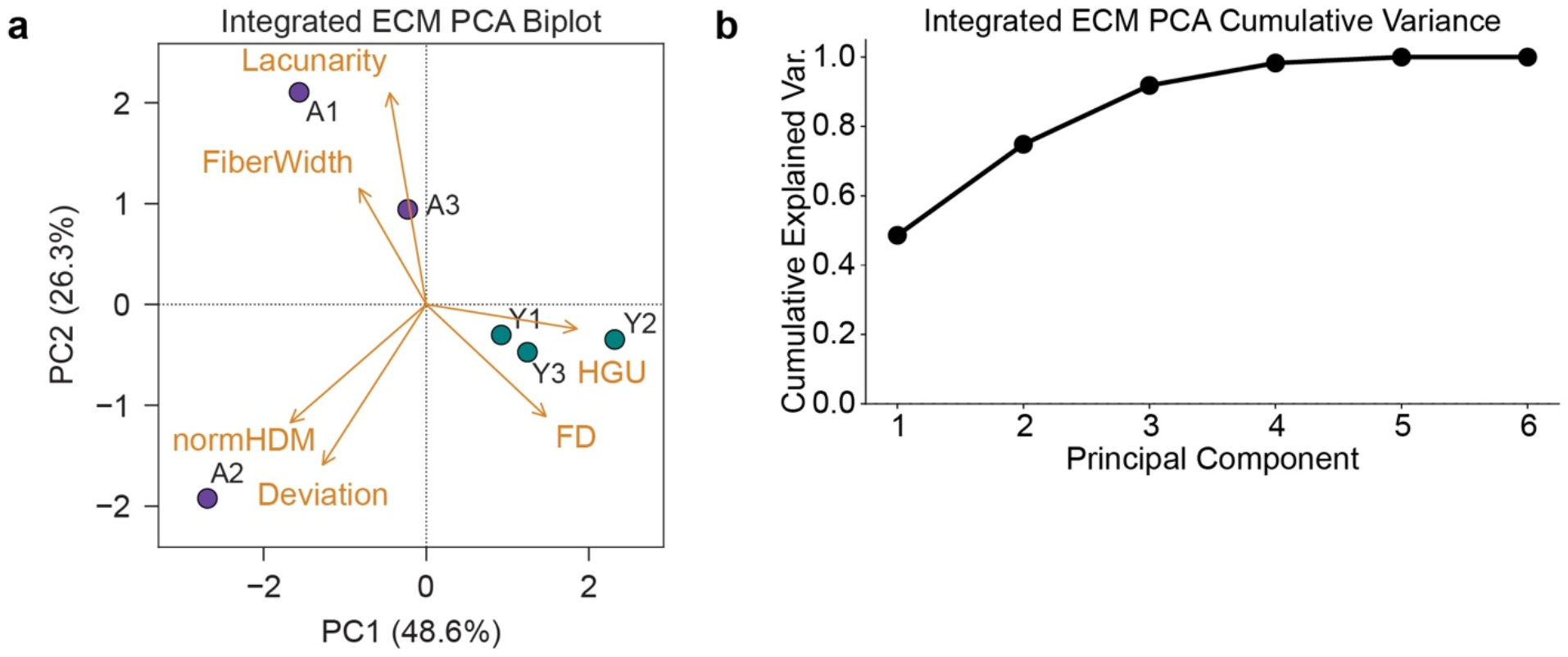
Validation of the integrated principal component analysis underlying the Architecture Index. (**a**) Principal component analysis (PCA) biplot of integrated ECM architectural features from individual young (Y1–Y3) and aged (A1–A3) biological replicates. Loading vectors indicate the contributions of high-density matrix (HDM), fractal dimension (FD), hyphal growth unit (HGU), lacunarity, deviation, and fiber width to the principal components.(**b**) Cumulative explained variance of the integrated PCA across the six principal components, demonstrating that PC1 accounted for the largest proportion of variance.

**Figure S3.**
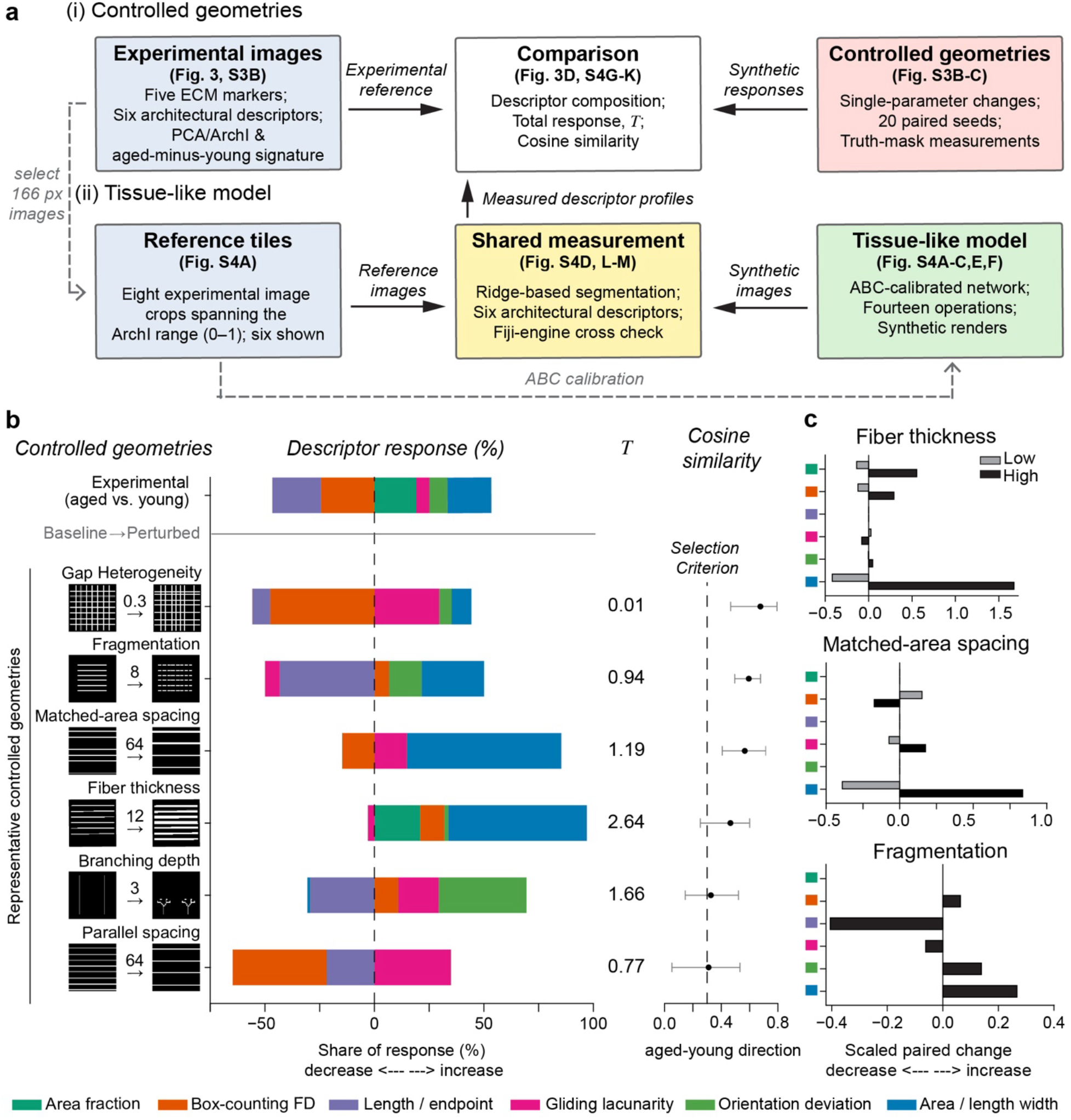
Synthetic ECM modeling workflow and descriptor responses to controlled geometric perturbations. **(a)** Overview of the synthetic simulation workflow using controlled geometries and a tissue-like model. Solid black arrows indicate the flow of experimental and synthetic data through the measurement and comparison procedures. Dashed gray arrows indicate the selection of experimental reference image crops and the calibration of the tissue-like model using reference-tile descriptor profiles through approximate Bayesian computation (ABC). **(b)** Controlled geometric perturbations and their agreement with the experimental aging signature. From left to right, the four columns show *Single perturbation*, showing representative baseline and perturbed truth masks from paired seed 0, with higher-endpoint parameter values indicated above the arrows; Descriptor response, showing the signed relative contributions (%) of the six architectural descriptors. For each controlled perturbation, median paired descriptor changes across 20 seeds were scaled by their respective 5th–95th percentile ranges and normalized by the sum of the six absolute scaled changes to obtain signed percentage contributions; (%); *Total response*, representing the total scaled response magnitude (T) represents the overall magnitude of the six scaled descriptor changes, calculated as ∑^6^ |*q_m_*|, where (*q_m_*) denotes the scaled median paired change for descriptor (m). Larger values indicate a greater collective descriptor response, irrespective of the direction of individual changes; and *Cosine similarity*, showing directional agreement between synthetic descriptor responses and the experimental aged-minus-young direction. Points indicate cosine similarity across all six hearts, whiskers show leave-one-heart-out ranges, and the dashed line marks the 0.30 selection criterion. **(c)** Individual descriptor responses to lower and higher geometric perturbation endpoints. Fiber thickness was varied from 4 px (baseline) to 2 and 12 px, while matched-area spacing was varied from 32 px to 18 and 64 px, with fiber width adjusted to maintain constant foreground area. Fragmentation divided each fiber from one continuous segment into eight separated pieces while maintaining constant foreground pixel count. Bars represent median paired descriptor changes across 20 seeds, scaled by each descriptor’s 5^th^–95^th^ percentile range. Light gray and dark bars indicate lower and higher endpoints, respectively; fragmentation includes only the higher endpoint. Positive and negative values indicate increases and decreases relative to baseline. Descriptor colors follow (b).

**Figure S4.**
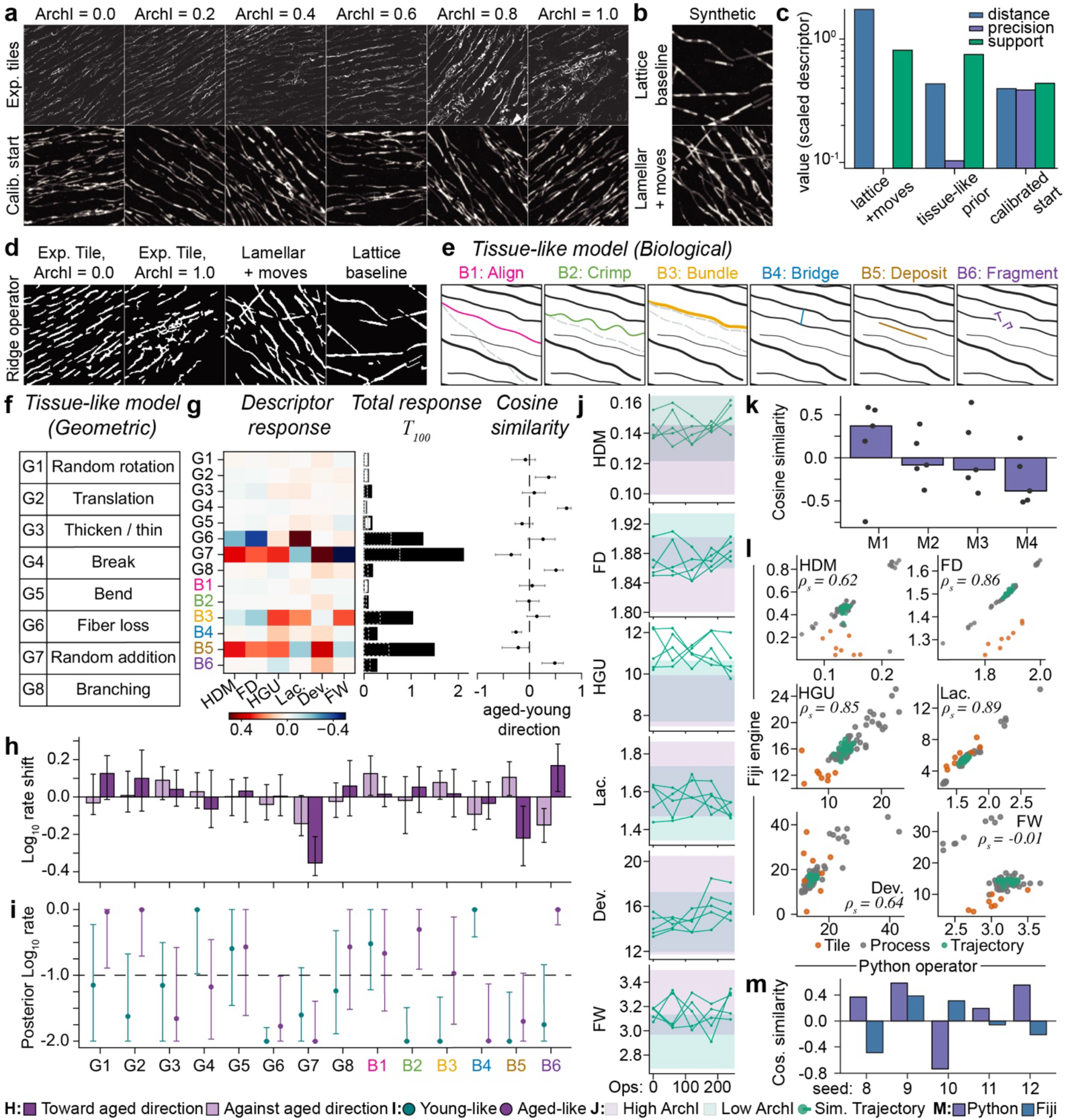
Tissue-like ECM network generation and descriptor responses to remodeling. **(a)** Representative experimental reference tiles and calibrated tissue-like networks. Six of the eight experimental reference tiles used for approximate Bayesian computation (ABC) calibration are shown, ordered by ArchI (0–1), alongside six fluorescence-like synthetic crops generated from the calibrated lamellar starting network. Synthetic crops represent the calibrated starting architecture and are not individually matched to the experimental tiles above. **(b)** Representative synthetic images showing a lattice baseline and a lamellar network following simulated remodeling operations (lamellar + moves), illustrating the different synthetic architectures examined in the modeling study. **(c)** Quantitative assessment of synthetic network realism. Three modeling approaches were compared against eight experimental reference tiles in scaled six-descriptor space: lattice with geometric moves, uncalibrated tissue-like networks (prior), and the calibrated tissue-like starting network. Metrics include median nearest-neighbor distance (blue), measuring proximity to experimental descriptor profiles; joint precision (purple), representing the fraction of synthetic networks with all six descriptors within the experimental ranges; and support (green), representing the fraction of experimental descriptor values within the synthetic 5^th^–95^th^ percentile intervals. Values are displayed on a logarithmic scale. **(d)** Shared ridge-based segmentation of experimental and synthetic ECM images. The experimental reference tiles (ArchI = 0.0 and 1.0) and synthetic examples shown in (A) and (B), respectively, were processed using the same ridge-based operator to generate binary masks. **(e)** Six biological remodeling operations (B1–B6) applied to a tissue-like lamellar network. Gray dashed lines indicate original fiber positions, colored lines indicate modified or newly added fibers, and black lines represent unchanged fibers. **(f)** Eight geometric remodeling operations (G1–G8) implemented in the tissue-like model, with perturbation parameters sampled from predefined distributions. **(g)** Individual tissue-like remodeling operations and their agreement with the experimental aging signature. From left to right, the three columns show *Descriptor response*, representing the median scaled changes in six architectural descriptors per 100 operations across 30 seeds. Responses were scaled by each descriptor’s 5^th^–95^th^ percentile range of the pooled measurements, with red and blue indicating increases and decreases, respectively; *Total response*, representing the combined magnitude of the six absolute scaled descriptor changes per 100 operations (T₁₀₀). Solid bars indicate observed responses, dashed gray outlines indicate the 95^th^ percentile of the sign-flip null distribution, and hollow bars indicate responses that did not exceed this threshold; and *Cosine similarity*, showing directional agreement between individual remodeling responses and the experimental aged-minus-young direction. Points indicate cosine similarities calculated using all six hearts, whiskers show leave-one-heart-out ranges, and the dashed vertical line marks zero similarity. **(h)** Remodeling-operation rate shifts associated with directional agreement with experimental aging. Bars represent median log₁₀ rate shifts relative to the prior median for mixtures moving toward (cosine ≥ 0.5, n = 170) or against (cosine ≤ −0.5, n = 257) the experimental aged-minus-young direction. Whiskers indicate 5th–95th percentile bootstrap intervals. **(i)** ABC-inferred remodeling-operation rates for young-like and aged-like reference architectures. Posterior log₁₀ rates were estimated using the mean six-descriptor profiles of the four lowest- and four highest-ArchI reference tiles through rejection ABC with regression adjustment. Points represent posterior medians, whiskers indicate 5^th^–95^th^ percentile intervals from the accepted 2% of draws, and the dashed horizontal line indicates the prior median (−1). Estimates are descriptive given the four reference tiles per group. **(j)** Architectural descriptor trajectories during tissue-like network remodeling using the selected mixture (M1, see K). The six descriptors were tracked over 240 remodeling operations across five simulation seeds, with measurements obtained from four fixed 166 × 166-pixel crops per checkpoint. Lines represent individual simulation trajectories, while shaded regions indicate the descriptor ranges of the four lowest-ArchI and four highest-ArchI experimental reference tiles. **(k)** Comparison of four candidate remodeling mixtures (M1–M4). Bars represent median cosine similarity to the experimental aged-minus-young direction across five simulation seeds, with points indicating individual seeds. M1 was selected for the remodeling trajectories in (J) based on the highest median cosine similarity. The operation probabilities for each mixture were: M1 (B6: 35%, B3: 25%, G1: 15%, G3: 15%, G6: 10%); M2 (B6: 70%, G1: 15%, G3: 15%); M3 (B6: 40%, G1: 20%, G2: 15%, G3: 15%, G6: 10%); and M4 (B6: 45%, G3: 25%, G1: 15%, B3: 15%). Operation codes are defined in (E) and (F). **(l)** Cross-check of architectural descriptor measurements between Python and Fiji-based routines across 108 experimental and synthetic images. Scatterplots compare Python-derived (X-axis) and Fiji-derived (Y-axis) measurements for the six descriptors. Points represent experimental reference tiles, simulated process endpoints, and remodeling trajectories. Fiji-based measurements used AnaMorf for FD, HGU, and lacunarity, and ImageJ-based routines for fiber deviation and width. Spearman rank correlation coefficients (ρₛ) are indicated in each plot. **(m)** Cross-check of remodeling trajectory agreement with the experimental aged-minus-young direction using Python and Fiji-based measurement routines. Bars indicate cosine similarities calculated from the six-descriptor changes of five simulation seeds (8–12), measured using the Python operator (purple) and Fiji-based routines (blue).

**Figure S5.**
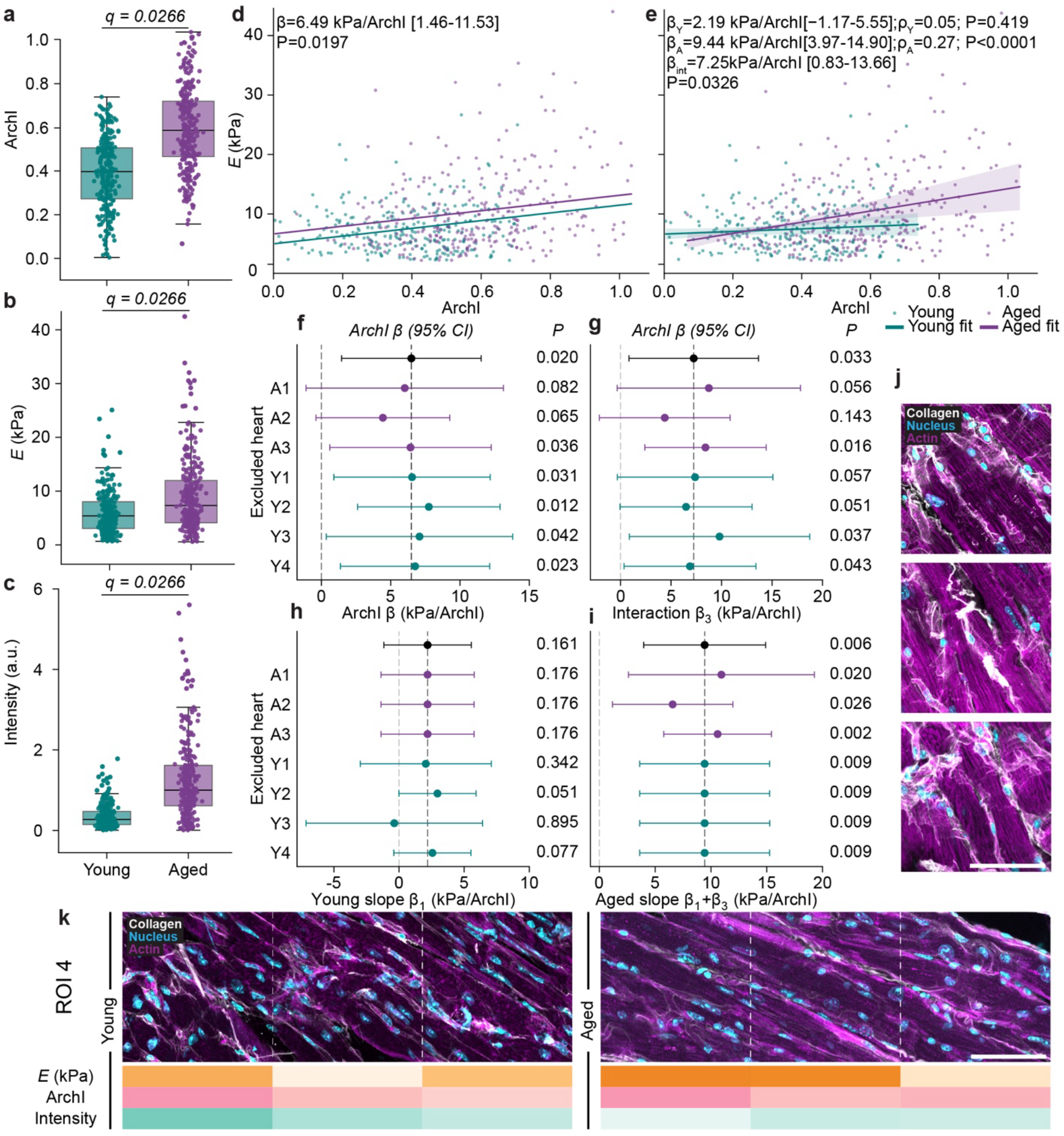
ECM architecture–stiffness associations and their age dependence in young and aged myocardium. **(a-c)** Distributions of (a) Architecture Index (ArchI), (b) apparent Young’s modulus, *E* (kPa), and (c) nuclei-normalized collagen integrated fluorescence intensity (IntDen/Nuc) in young and aged myocardium. Points represent individual spatial measurements. Boxes indicate the interquartile range (IQR), central lines indicate the median, and whiskers extend to the most extreme observations within 1.5 × IQR of the corresponding quartile. Age-group comparisons were assessed using ordinary least squares (OLS) regression with CR1 cluster-robust standard errors. Unadjusted *P* values and false discovery rate (FDR)-adjusted *q* values are indicated. **(d)** Age-adjusted association between ArchI and *E*, showing parallel fitted regression lines with a common ArchI slope. **(e)** ArchI × age interaction model showing age-specific fitted regression lines for young and aged myocardium. Shaded regions represent pointwise 95% confidence intervals for the fitted mean, calculated using the full CR1 cluster-robust covariance matrix with biological replicate (heart) as the clustering variable. Fitted lines and confidence intervals are restricted to the observed ArchI range of each age group. Spearman’s rank correlation coefficients (ρ) and corresponding *P* values are indicated for each age group. **(f)** Leave-one-heart-out (LOHO) sensitivity analysis of the age-adjusted ArchI regression coefficient. **(g–i)** LOHO sensitivity analyses of (G) the ArchI × age interaction coefficient (β₃), (H) the young slope (β₁), and (I) the aged slope (β₁ + β₃). Points represent coefficient estimates, and horizontal bars indicate 95% confidence intervals. Black indicates the full-dataset estimate, while teal and purple indicate exclusion of young and aged hearts, respectively. Dashed vertical lines indicate zero, where shown. **(j)** Three representative confocal images of myocardial regions with ArchI values marginally exceeding 1.0, selected from five observations (0.91% of total images analysed), all from aged hearts. **(k)** Two additional representative regions of interest (ROIs) from independent young and aged hearts, comprising one additional heart per age group, illustrating the ECM architectures associated with Fig. 5. Regression analyses included 548 spatially matched measurements from seven hearts (four young and three aged). Standard errors were clustered by heart replicates, with two-sided *t*-based inference using six degrees of freedom for full-dataset analyses and five for LOHO analyses. Regression *P* values were unadjusted. Teal and purple denote young and aged groups, respectively. Detailed statistical results are provided in **Supplementary Table S11**.

## Supplementary tables

**Supplementary Table 1:**
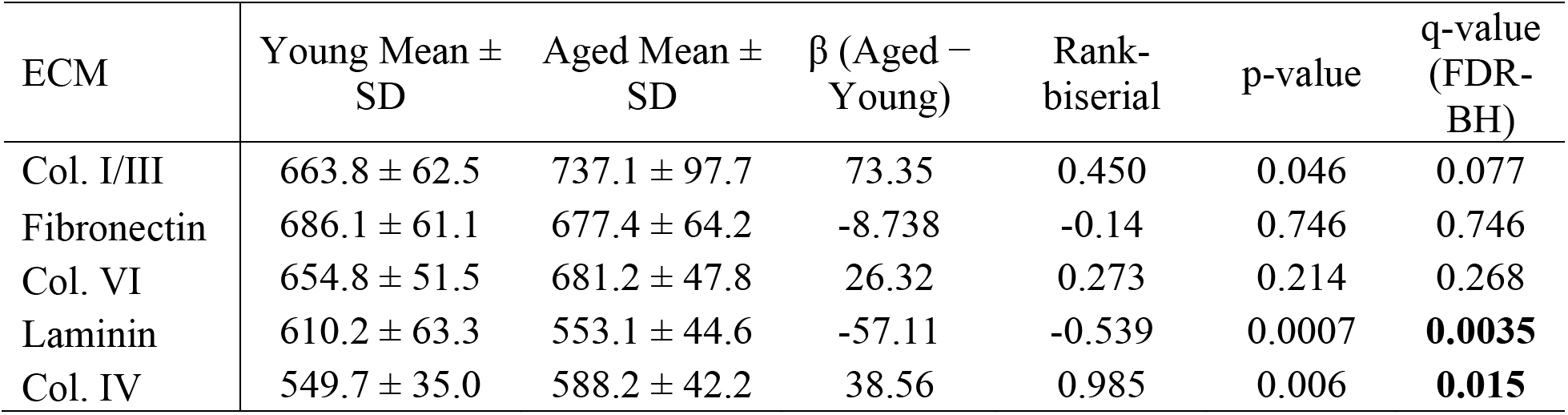
OLS-CR1 regression results for area-normalized integrated density (Figure 1b). *P* values were adjusted across the five ECM components using the Benjamini–Hochberg false discovery rate (FDR) procedure, with *q* < 0.05 considered statistically significant.

| ECM | Young Mean ± SD | Aged Mean ± SD | β (Aged − Young) | Rank-biserial | p-value | q-value (FDR-BH) |
|---|---|---|---|---|---|---|
| Col. I/III | 663.8 ± 62.5 | 737.1 ± 97.7 | 73.35 | 0.450 | 0.046 | 0.077 |
| Fibronectin | 686.1 ± 61.1 | 677.4 ± 64.2 | -8.738 | -0.14 | 0.746 | 0.746 |
| Col. VI | 654.8 ± 51.5 | 681.2 ± 47.8 | 26.32 | 0.273 | 0.214 | 0.268 |
| Laminin | 610.2 ± 63.3 | 553.1 ± 44.6 | -57.11 | -0.539 | 0.0007 | **0.0035** |
| Col. IV | 549.7 ± 35.0 | 588.2 ± 42.2 | 38.56 | 0.985 | 0.006 | **0.015** |

**Supplementary Table 2:** OLS-CR1 regression results for Architecture Index (ArchI), (Figure 2b & d). Component-specific *P* values were adjusted for multiple comparisons using the Benjamini–Hochberg false discovery rate (FDR) procedure, with *q* < 0.05 considered statistically significant.

| ECM | Young Mean ± SD | Aged Mean ± SD | β (Aged − Young) | Rank-biserial | p-value | q-value (FDR-BH) |
|---|---|---|---|---|---|---|
| Overall | 0.42 ± 0.16 | 0.55 ± 0.16 | 0.128 | 0.243 | **0.0056** | - |
| Col. I/III | 0.41 ± 0.17 | 0.57 ± 0.17 | 0.163 | 0.281 | 0.0059 | **0.015** |
| Fibronectin | 0.38 ± 0.13 | 0.57 ± 0.11 | 0.188 | 0.443 | 0.0137 | **0.023** |
| Col. VI | 0.51 ± 0.13 | 0.49 ± 0.12 | -0.021 | -0.027 | 0.4886 | 0.611 |
| Laminin | 0.43 ± 0.16 | 0.57 ± 0.13 | 0.138 | 0.247 | 0.0058 | **0.015** |
| Col. IV | 0.49 ± 0.14 | 0.50 ± 0.19 | 0.008 | -0.026 | 0.9018 | 0.902 |

**Supplementary Table 3.**
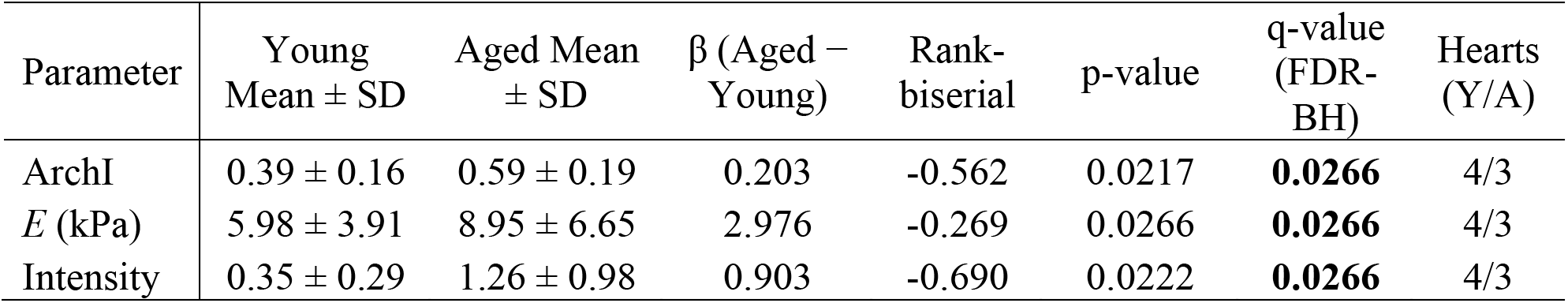
OLS-CR1 regression results for Architecture Index (ArchI), apparent Young’s modulus, and nuclei-normalized collagen I/III fluorescence intensity (Figures 5 and S5). Collagen fluorescence intensity was quantified as raw integrated density normalized to the number of nuclei within each spatially matched region (RawIntDen/Nuc). *P* values were adjusted across the three outcomes using the Benjamini–Hochberg false discovery rate (FDR) procedure.

**Supplementary Table 4:** Principal component loadings from principal component analysis (PCA) used to derive the Architecture Index. PC1 loadings were used as weighting coefficients to calculate the Architecture Index, whereas the remaining principal components are provided for completeness. HDM, high-density matrix; FD, fractal dimension; HGU, hyphal growth unit; Fiber deviation, length-weighted angular deviation of fibers; Fiber width, mean fiber width.

| PCA loadings |  |  |  |  |  |  |
|---|---|---|---|---|---|---|
| Component | HDM | FD | HGU | Lacunarity | Deviation | Fiber Width |
| PC1 | -0.506 | 0.446 | 0.562 | -0.136 | -0.385 | -0.251 |
| PC2 | -0.355 | -0.337 | -0.073 | 0.633 | -0.481 | 0.350 |
| PC3 | 0.234 | 0.117 | 0.087 | -0.484 | -0.362 | 0.747 |
| PC4 | -0.021 | 0.750 | -0.124 | 0.435 | 0.335 | 0.348 |
| PC5 | -0.005 | -0.311 | 0.745 | 0.134 | 0.496 | 0.291 |
| PC6 | 0.750 | 0.124 | 0.318 | 0.373 | -0.362 | -0.225 |

**Supplementary Table 5:** Explained variance ratio and cumulative variance for principal component analysis (PCA) of individual extracellular matrix components and the combined dataset.

| **Col. I/III** |  |  | **Laminin** |  |  |
|---|---|---|---|---|---|
| Component | Explained Variance Ratio | Cumulative Variance | Component | Explained Variance Ratio | Cumulative Variance |
| PC1 | 0.560 | 0.560 | PC1 | 0.545 | 0.545 |
| PC2 | 0.318 | 0.878 | PC2 | 0.318 | 0.863 |
| PC3 | 0.089 | 0.968 | PC3 | 0.106 | 0.969 |
| PC4 | 0.032 | 1.000 | PC4 | 0.023 | 0.992 |
| PC5 | 0.000 | 1.000 | PC5 | 0.009 | 1.000 |
| PC6 | 0.000 | 1.000 | PC6 | 0.000 | 1.000 |
| **Fibronectin** |  |  | **Col. IV** |  |  |
| Component | Explained Variance Ratio | Cumulative Variance | Component | Explained Variance Ratio | Cumulative Variance |
| PC1 | 0.621 | 0.621 | PC1 | 0.727 | 0.727 |
| PC2 | 0.273 | 0.893 | PC2 | 0.127 | 0.854 |
| PC3 | 0.082 | 0.975 | PC3 | 0.119 | 0.973 |
| PC4 | 0.024 | 0.999 | PC4 | 0.022 | 0.995 |
| PC5 | 0.001 | 1.000 | PC5 | 0.005 | 1.000 |
| PC6 | 0.000 | 1.000 | PC6 | 0.000 | 1.000 |
| **Col. VI** |  |  | **Combined** |  |  |
| Component | Explained Variance Ratio | Cumulative Variance | Component | Explained Variance Ratio | Cumulative Variance |
| PC1 | 0.521 | 0.521 | PC1 | 0.486 | 0.486 |
| PC2 | 0.286 | 0.807 | PC2 | 0.263 | 0.749 |
| PC3 | 0.154 | 0.961 | PC3 | 0.170 | 0.918 |
| PC4 | 0.030 | 0.991 | PC4 | 0.065 | 0.983 |
| PC5 | 0.009 | 1.000 | PC5 | 0.017 | 1.000 |
| PC6 | 0.000 | 1.000 | PC6 | 0.000 | 1.000 |

**Supplementary Table 6:** ECM geometric descriptors and ECM compartment classifications used for quantitative analysis of cardiac extracellular matrix remodeling.

| **Geometric Feature** | Quantification | Interpretation in myocardial ECM |
|---|---|---|
| **High Density Matrix (HDM):**<br>Matrix occupancy | Proportion of pixels classified as ECM based on an intensity threshold. | Higher HDM reflects greater local ECM occupancy and may represent increased matrix deposition or compaction, whereas lower HDM reflects a sparse ECM network. Increased collagen density has similarly been associated with more tightly packed myocardial fibrosis [63]. |
| **Fractal Dimension (FD):**<br>Space-filling organization | Calculated using box-counting fractal dimension method. | Higher FD indicates greater space-filling across scales, whereas lower FD indicates less space-filling organization. In myocardial fibrosis, lower FD with higher collagen density has been associated with tightly packed reactive fibrosis, whereas higher FD was associated with more space-filling reparative fibrosis [63]. |
| **Lacunarity:**<br>Spatial heterogeneity | Quantifies variation in pixel intensity across different window sizes. | Higher lacunarity reflects greater variation in matrix-free gaps and therefore a more spatially heterogeneous myocardial ECM, whereas lower lacunarity indicates a more uniformly distributed matrix network [19,64]**.** |
| **HGU (Hyphal Growth Unit):**<br>Network continuity | The total fiber length per fiber end point. | Higher HGU indicates greater fiber length per endpoint and is consistent with a more continuous matrix network, whereas lower HGU reflects greater discontinuity. During fibrotic ECM remodeling, increased fiber endpoints accompanied by reduced HGU have been reported, consistent with greater network fragmentation [19,65]. |
| **Deviation:**<br>Fiber orientation | Deviation of fibers from the dominant matrix orientation, using a major-axis-weighted axial mean. | Higher fiber deviation reflects greater angular dispersion from the preferential matrix orientation, whereas lower deviation reflects stronger directional organization. This is relevant because myocardial ECM is preferentially organized with the underlying myocardial architecture [66,67]. |
| **Fiber Width:**<br>Matrix fiber thickness | Minor-axis length of segmented ECM fibers, weighted by major-axis length. | Higher values indicate thicker or more bundled matrix structures, whereas lower values indicate finer structures. Apparent width is component dependent and may reflect bundling, or local matrix aggregation [68]. |

**Supplementary Table 7:** Age-associated changes in extracellular matrix architectural descriptors across individual matrix components. Regression coefficients (β), unadjusted P values (P), and Benjamini–Hochberg false discovery rate (FDR)-adjusted P values (q) are reported for each architectural descriptor, comparing aged with young myocardium. Positive and negative β values indicate increases and decreases, respectively, in aged myocardium relative to young myocardium. Asterisks (*) indicate statistically significant differences after FDR correction (q < 0.05). HDM, high-density matrix; FD, fractal dimension; Lac., lacunarity; HGU, hyphal growth unit; Dev., deviation; FW, fiber width.

| **ECM**<br>β (p, q) | **HDM<br>(x10<sup>-3</sup>)** | **FD** | **Lac.** | **HGU** | **Dev.** | **FW** |
|---|---|---|---|---|---|---|
| Fibrillar Collagen | 0.852; 0.009; 0.018 * | -0.0409; 0.003; 0.017 * | -1.59; 0.675; 0.675 | -16.7; 0.053; 0.079 | 0.609; 0.386; 0.463 | 0.115; 0.007; 0.018 * |
| Fibro-nectin | -0.201; 0.541; 0.650 | -0.158; 0.117; 0.350 | 41.3; 0.191; 0.354 | -28.7; 0.236; 0.354 | 4.95; 0.012; 0.070 | 0.0231; 0.731; 0.731 |
| Collagen Type VI | 0.181; 0.369; 0.554 | -0.0436; 0.210; 0.554 | -8.91; 0.599; 0.718 | -0.216; 0.957; 0.957 | -2.06; 0.002; 0.014 * | -0.0215; 0.357; 0.554 |
| Laminin | 0.325; 0.619; 0.742 | -0.0697; 0.050; 0.134 | -0.287; 0.972; 0.972 | -34.2; 0.004; 0.025 * | 2.08; 0.089; 0.134 | -0.0782; 0.082; 0.134 |
| Collagen Type IV | 1.34; 0.160; 0.240 | 0.0318; 0.450; 0.450 | -51.7; 0.088; 0.240 | 6.1; 0.279; 0.335 | -2.61; 0.154; 0.240 | 0.108; 0.013; 0.080 |

**Supplementary Table 8:** Antibodies and dyes used for IHC

| **Target** | **Host species/ Product** | **Vendor & Catalog #** | **Dilution** |
|---|---|---|---|
| Fibrillar Collagen | Recombinant EGFP–CNA35 fusion protein | Addgene plasmid #61603 | 1 uM |
| Fibronectin | Rabbit, polyclonal IgG | Abcam, ab2413 | 1:100 |
| Collagen Type VI | Rabbit, polyclonal IgG | Novus Biologicals, NB120-6588 | 1:50 |
| Laminin α2 | Rat, monoclonal IgG1 | Sigma-Aldrich, L0663 | 1:100 |
| Collagen Type IV | Rabbit, polyclonal IgG | Abcam, ab6586 | 1:300 |
| F-actin | CF® Phalloidin | Biotium, 00044 | 1:250 |
| Nuclei | Hoechst 33342 | Invitrogen, H3570 | 1:2000 |
| **Secondary Antibodies** |  |  |  |
| Rabbit IgG | Goat anti-rabbit IgG (H+L), highly cross-adsorbed, CF®640R | Biotium, 20176 | 1:1000 |
| Rat IgG | Goat anti-rat IgG (H+L), highly cross-adsorbed, CF®647 | Biotium, 20283 | 1:1000 |

**Supplementary Table 9:** TWOMBLI analysis parameters. Parameter settings used for the automated processing and quantification of extracellular matrix (ECM) architecture from confocal fluorescence images using TWOMBLI in Fiji/ImageJ. The same parameter settings were applied across young and aged cardiac tissue samples.

| **Parameter** | **Value** |
|---|---|
| Contrast saturation | 0.35 |
| Minimum line width | 5 |
| Maximum line width | 20 |
| Minimum curvature window | 10 |
| Maximum curvature window | 10 |
| Minimum branch length | 40 |
| Maximum display HDM | 245 |

**Supplementary Table 10:** Heart-level descriptive statistics for ECM-specific Architecture Index (ArchI). Individual heart means (Y1–Y3 and A1–A3), group means, and 95% confidence intervals (CIs) are reported for each ECM component and the integrated ArchI (n = 3 hearts per age group). Group means and CIs were calculated from independent heart means using Student’s t-distribution (df = 2). Corresponding image-level descriptive statistics and OLS-CR1 regression results are provided in Supplementary Table 2.

| ECM component | Group | Group mean (95%CI) | Individual heart means (n =3) |  |  |
|---|---|---|---|---|---|
| Fibrillar collagen | Young | 0.408 (0.352–0.464) | 0.433 | 0.401 | 0.390 |
|  | Aged | 0.572 (0.422–0.722) | 0.602 | 0.611 | 0.502 |
| Fibronectin | Young | 0.385 (0.164–0.606) | 0.337 | 0.331 | 0.488 |
|  | Aged | 0.56 (0.461–0.658) | 0.553 | 0.602 | 0.524 |
| Collagen VI | Young | 0.507 (0.390–0.624) | 0.470 | 0.490 | 0.560 |
|  | Aged | 0.484 (0.425–0.542) | 0.475 | 0.465 | 0.510 |
| Laminin | Young | 0.434 (0.375–0.492) | 0.451 | 0.407 | 0.443 |
|  | Aged | 0.57 (0.455–0.686) | 0.581 | 0.61 | 0.519 |
| Collagen IV | Young | 0.516 (0.253–0.779) | 0.637 | 0.467 | 0.443 |
|  | Aged | 0.507 (0.287–0.727) | 0.605 | 0.486 | 0.431 |
| Overall | Young | 0.427 (0.382–0.471) | 0.443 | 0.408 | 0.428 |
|  | Aged | 0.555 (0.439–0.671) | 0.578 | 0.586 | 0.501 |

**Supplementary Table 11:**
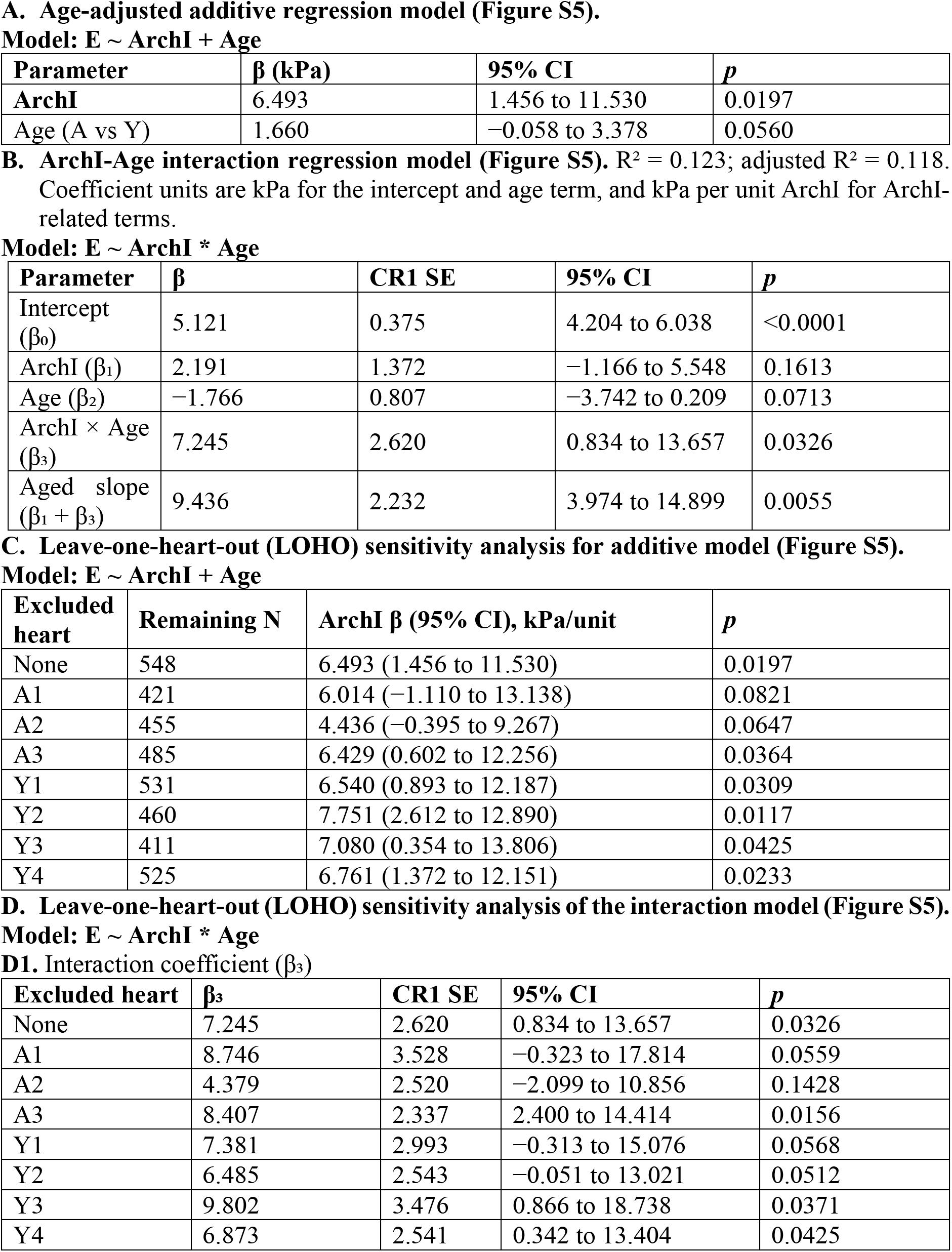
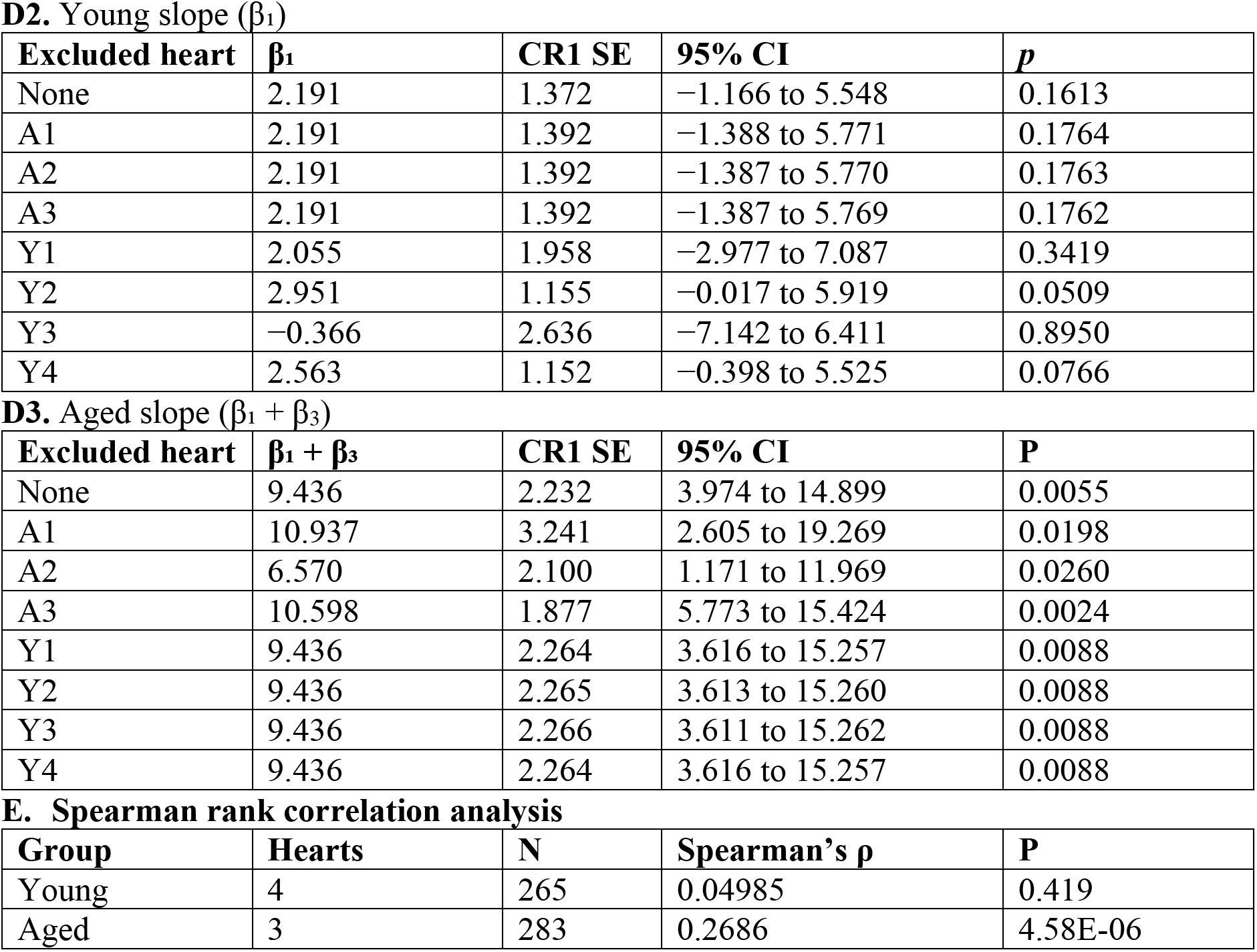
Regression statistics and leave-one-heart-out sensitivity analyses of spatially matched ECM architecture–stiffness associations.

| **A. Age-adjusted additive regression model (Figure S5).** |  |  |  |  |
|---|---|---|---|---|
| **Model: E ~ ArchI + Age** |  |  |  |  |
| **Parameter** | **β (kPa)** | **95% CI** | ***p*** |  |
| **ArchI** | 6.493 | 1.456 to 11.530 | 0.0197 |  |
| Age (A vs Y) | 1.660 | −0.058 to 3.378 | 0.0560 |  |
| **B. ArchI-Age interaction regression model (Figure S5).** R<sup>2</sup> = 0.123; adjusted R<sup>2</sup> = 0.118. Coefficient units are kPa for the intercept and age term, and kPa per unit ArchI for ArchI-related terms. |  |  |  |  |
| **Model: E ~ ArchI \* Age** |  |  |  |  |
| **Parameter** | **β** | **CR1 SE** | **95% CI** | ***p*** |
| Intercept (β<sub>0</sub>) | 5.121 | 0.375 | 4.204 to 6.038 | <0.0001 |
| ArchI (β<sub>1</sub>) | 2.191 | 1.372 | −1.166 to 5.548 | 0.1613 |
| Age (β<sub>2</sub>) | −1.766 | 0.807 | −3.742 to 0.209 | 0.0713 |
| ArchI × Age (β<sub>3</sub>) | 7.245 | 2.620 | 0.834 to 13.657 | 0.0326 |
| Aged slope (β<sub>1</sub> + β<sub>3</sub>) | 9.436 | 2.232 | 3.974 to 14.899 | 0.0055 |
| **C. Leave-one-heart-out (LOHO) sensitivity analysis for additive model (Figure S5).** |  |  |  |  |
| **Model: E ~ ArchI + Age** |  |  |  |  |
| **Excluded heart** | **Remaining N** | **ArchI β (95% CI), kPa/unit** | ***p*** |  |
| None | 548 | 6.493 (1.456 to 11.530) | 0.0197 |  |
| A1 | 421 | 6.014 (−1.110 to 13.138) | 0.0821 |  |
| A2 | 455 | 4.436 (−0.395 to 9.267) | 0.0647 |  |
| A3 | 485 | 6.429 (0.602 to 12.256) | 0.0364 |  |
| Y1 | 531 | 6.540 (0.893 to 12.187) | 0.0309 |  |
| Y2 | 460 | 7.751 (2.612 to 12.890) | 0.0117 |  |
| Y3 | 411 | 7.080 (0.354 to 13.806) | 0.0425 |  |
| Y4 | 525 | 6.761 (1.372 to 12.151) | 0.0233 |  |
| **D. Leave-one-heart-out (LOHO) sensitivity analysis of the interaction model (Figure S5).** |  |  |  |  |
| **Model: E ~ ArchI \* Age** |  |  |  |  |
| **D1.** Interaction coefficient (β<sub>3</sub>) |  |  |  |  |
| **Excluded heart** | **β<sub>3</sub>** | **CR1 SE** | **95% CI** | ***p*** |
| None | 7.245 | 2.620 | 0.834 to 13.657 | 0.0326 |
| A1 | 8.746 | 3.528 | −0.323 to 17.814 | 0.0559 |
| A2 | 4.379 | 2.520 | −2.099 to 10.856 | 0.1428 |
| A3 | 8.407 | 2.337 | 2.400 to 14.414 | 0.0156 |
| Y1 | 7.381 | 2.993 | −0.313 to 15.076 | 0.0568 |
| Y2 | 6.485 | 2.543 | −0.051 to 13.021 | 0.0512 |
| Y3 | 9.802 | 3.476 | 0.866 to 18.738 | 0.0371 |
| Y4 | 6.873 | 2.541 | 0.342 to 13.404 | 0.0425 |
| **D2.** Young slope (β<sub>1</sub>) |  |  |  |  |
| **Excluded heart** | **β<sub>1</sub>** | **CR1 SE** | **95% CI** | ***p*** |
| None | 2.191 | 1.372 | −1.166 to 5.548 | 0.1613 |
| A1 | 2.191 | 1.392 | −1.388 to 5.771 | 0.1764 |
| A2 | 2.191 | 1.392 | −1.387 to 5.770 | 0.1763 |
| A3 | 2.191 | 1.392 | −1.387 to 5.769 | 0.1762 |
| Y1 | 2.055 | 1.958 | −2.977 to 7.087 | 0.3419 |
| Y2 | 2.951 | 1.155 | −0.017 to 5.919 | 0.0509 |
| Y3 | −0.366 | 2.636 | −7.142 to 6.411 | 0.8950 |
| Y4 | 2.563 | 1.152 | −0.398 to 5.525 | 0.0766 |
| **D3.** Aged slope (β<sub>1</sub> + β<sub>3</sub>) |  |  |  |  |
| **Excluded heart** | **β<sub>1</sub> + β<sub>3</sub>** | **CR1 SE** | **95% CI** | **P** |
| None | 9.436 | 2.232 | 3.974 to 14.899 | 0.0055 |
| A1 | 10.937 | 3.241 | 2.605 to 19.269 | 0.0198 |
| A2 | 6.570 | 2.100 | 1.171 to 11.969 | 0.0260 |
| A3 | 10.598 | 1.877 | 5.773 to 15.424 | 0.0024 |
| Y1 | 9.436 | 2.264 | 3.616 to 15.257 | 0.0088 |
| Y2 | 9.436 | 2.265 | 3.613 to 15.260 | 0.0088 |
| Y3 | 9.436 | 2.266 | 3.611 to 15.262 | 0.0088 |
| Y4 | 9.436 | 2.264 | 3.616 to 15.257 | 0.0088 |
| **E. Spearman rank correlation analysis** |  |  |  |  |
| **Group** | **Hearts** | **N** | **Spearman’s ρ** | **P** |
| Young | 4 | 265 | 0.04985 | 0.419 |
| Aged | 3 | 283 | 0.2686 | 4.58E-06 |

**Supplementary Table 12:** Biological replicates and measurement counts used for ECM architecture and correlative structure–stiffness analyses. Individual mice represent independent biological replicates (N), reported as Young/Aged (Y/A). Within-heart measurement counts (n) are listed for individual hearts, followed by the pooled total (P) for each age group. Individual counts follow the order Y1–Y3/A1–A3 for ECM architectural analyses and Y1–Y4/A1–A3 for correlative structure–stiffness analyses. For integrated ArchI derivation, image-level measurements were aggregated at the heart level before PCA. Correlative measurements represent individual spatially matched positions or registration grids, with grid-level means used for Fig. 5B. Measurement units and corresponding figures are indicated.

| Analysis | Hearts (N)<br>Y/A | Young (n)<br>N/P | Aged (n)<br>N/P | Measurement unit | Figure |
|---|---|---|---|---|---|
| **Architectural Analysis** |  |  |  |  |  |
| Fibrillar collagen | 3/3 | 68/99/93<br>260 | 76/125/121<br>322 | Images | 1 |
| Fibronectin | 3/3 | 9/21/11<br>41 | 15/23/19<br>57 |  |  |
| Collagen VI | 3/3 | 19/18/18<br>55 | 18/23/26<br>67 |  |  |
| Laminin | 3/3 | 15/23/22<br>60 | 15/22/22<br>59 |  |  |
| Collagen IV | 3/3 | 6/15/10<br>31 | 9/13/16<br>38 |  |  |
| **ArchI Derivation** |  |  |  |  |  |
| Integrated PCA & ArchI | 3/3 | 447 | 543 | Images (aggregated by Heart) | 2 |
| **Stiffness-Structure Analysis** |  |  |  |  |  |
| Matched spatial measurements | 4/3 | 17/88/137/23<br>265 | 127/93/63<br>283 | Images/Heart | 5 |
| Fig 5b: Scatter plot | 4/3 | 7/11/21/3<br>42 | 15/12/8<br>35 | Grid |  |

